# A NEW DYSTROPHIN DEFICIENT RAT MODEL MIRRORING EXON SKIPPING IN PATIENTS WITH DMD EXON 45 DELETIONS

**DOI:** 10.1101/2025.05.30.656540

**Authors:** Abbass Jaber, Tao Wang, Cynthia Daoud, Sonia Albini, Guillaume Corre, Jessica Bellec, Matteo Bovolenta, Alan Dorval, Auriane Dubois, Louise Philidet, Ganesh Warthi, Isabelle Richard

## Abstract

Mutations in the dystrophin (DMD) gene can cause a spectrum of muscle-wasting disorders ranging from the milder Becker muscular dystrophy (BMD) to the more severe Duchenne muscular dystrophy (DMD). Among these, exon 45 deletion is the most frequently reported single exon deletion in DMD patients worldwide. In this study, we generated a novel rat model with an exon 45 deletion using CRISPR/Cas9 technology. The *Dmd^Δ45^* rat recapitulate key clinical and molecular features of DMD, including progressive skeletal muscle degeneration, cardiac dysfunction, cognitive deficits, elevated circulating muscle damage biomarkers, impaired muscle function, and overall reduced lifespan. Transcriptomics analyses confirmed the deletion of exon 45 and revealed gene expression patterns consistent with dystrophin deficiency. In the skeletal muscle, RNA-seq profiles demonstrated a transition from early stress responses and regenerative activity at 6 months to chronic inflammation, fibrosis, and metabolic dysfunction by 12 months. Similarly, the cardiac transcriptomic shifted from an early inflammatory and stress-responsive state to one characterized by fibrotic remodelling and metabolic impairment. Despite these pathological features, the *Dmd^Δ45^* rats exhibited a milder phenotype than other DMD rat models. This attenuation may be attributed to spontaneous exon 44 skipping, which partially restores the reading frame and results in an age-dependent increase in revertant dystrophin-positive fibres. Further analysis indicated downregulation of spliceosome-related genes, suggesting a potential mechanism driving exon skipping in this model. In summary, the *Dmd^Δ45^* rat represents a valuable model for investigating both the molecular determinants of phenotypic variability and the endogenous mechanisms of exon skipping. These findings offer important insights for the development of personalized exon-skipping therapies, particularly for DMD patients with exon 45 deletions.

## INTRODUCTION

Duchenne muscular dystrophy (DMD) is a severe X-linked recessive muscular disorder caused by mutations in the *DMD* gene, which encodes dystrophin—a structural protein critical for muscle fibre integrity [1, 2]. With an incidence of approximately 1 in 5,000 male births, DMD is the most common and devastating form of childhood muscular dystrophy [3, 4]. Mutations in the same gene can also cause Becker muscular dystrophy (BMD), a clinically milder form characterized by the expression of partially functional dystrophin forms, and that typically presents with later onset and slower disease progression [5, 6].

The protein product of *DMD*, dystrophin, is predominantly expressed in skeletal and cardiac muscle tissues. It is a sub-sarcolemmal protein with several bindings domains, enabling it to establish a mechanical bridge between the extracellular matrix (ECM) and the cytoskeleton. Dystrophin is a pivotal constituent of the dystrophin-associated glycoprotein complex (DAGC) Ervasti, 1991 #88}, crucial not only for maintaining muscle fibre rigidity [7], but also for shielding the muscle against mechanical stress encountered during contractions [8, 9]. Its deficiency leads to perturbation of the assembly of the DAGC and destabilization the muscle membrane, making it more fragile and sensitive to mechanical stress [10, 11].

Clinically, DMD manifests early, usually under the age of 2-3 years, with proximal muscle weakness, particularly in the lower limbs, accompanied by elevated serum creatine kinase (CK) levels as a result of ongoing muscle fibre damage [3]. Muscle weakness progresses rapidly, leading to difficulties standing up and an inability to walk by 10-12 years of age. Respiratory muscles become affected, necessitating assisted ventilation at around 20 years old. While motor deficits dominate the early clinical picture, cognitive impairments, especially in working memory and executive function, are also frequently reported [12], although these typically remain stable over time. Cardiac and respiratory complications generally emerge during the second or third decade of life and are now recognized as the leading causes of mortality in DMD [13, 14].

The *DMD* gene itself is the largest known protein-coding gene, spanning over 2.6 million base pairs and comprising 79 exons. Owing to its size, it is highly susceptible to mutation, with thousands of pathogenic variants identified in DMD and BMD patients [15]. In DMD, these mutations typically result in the absence of functional dystrophin protein, with ∼60–70% being exon deletions and ∼20% comprising point mutations, small insertions, or deletions [15–17]. In contrast, BMD mutations tend to preserve the open reading frame, allowing the production of truncated but partially functional dystrophins. Mutation hotspots have been identified in the *DMD* gene, notably in exons 3–9 and exons 45–55 [18, 19], with the latter accounting for nearly half of all deletions observed in DMD patients worldwide.

Current DMD care relies on an early, multidisciplinary management focused on symptom relief with especially the use of glucocorticoids to slow disease progression. In parallel, therapeutic strategies aimed at restoring dystrophin expression have shown promise, with some gaining regulatory approval. These include exon skipping using antisense oligonucleotides (AONs) to bypass mutated exons during mRNA splicing [22]. Another promising, mutation-independent approach involves the delivery of shortened dystrophin constructs (micro-dystrophins) using adeno-associated viral (AAV) vectors [23].

The exon 45–55 region is of particular interest for exon skipping therapies, as restoring the reading frame here can yield a BMD-like dystrophin and milder clinical phenotype. Skipping exon 45, in particular, has shown therapeutic potential [24, 25], underscoring the importance of disease models that replicate this hotspot for the development and testing of AONs and gene editing tools. While the *mdx* mouse model, harbouring a point mutation in exon 23, is widely used, its mild phenotype and limited cardiac involvement reduce its translational relevance [26]. In contrast, DMD rat models, enabled by recent advances in genome-editing technologies, exhibit more severe and human-like disease progression, including pronounced muscle degeneration, fibrosis, and cardiac involvement [27–30].

To fill the gap in models targeting patient-relevant hotspots, we generated a novel *Dmd^Δ45^* rat model featuring a targeted deletion of exon 45. This model exhibited hallmark features of DMD, including progressive impairments in skeletal muscle, cardiac function, and cognitive performance, alongside elevated muscle damage biomarkers, impaired muscle function and overall reduced lifespan. Interestingly, despite these features, the phenotype in this model was less severe compared to other reported DMD rat models. This attenuation may be attributed in part to preferential progressive skipping of exon 44, which restores the reading frame and results in the expression of partially functional dystrophin in the form of revertant fibres. Transcriptomic analysis revealed age-dependent dysregulation of genes involved in fibrosis and the spliceosome, shedding light on disease mechanisms and the biological basis of spontaneous exon skipping. Collectively, the *Dmd^Δ45^* rat represents a valuable model for dissecting the pathophysiology of DMD/BMD associated with exon 45–55 mutations and offers a robust preclinical platform for the development and testing of exon-skipping and gene-editing therapies targeting this critical mutational hotspot.

## RESULTS

### Generation and validation of a *Dmd^Δ45^* rat model

To generate the *Dmd*^Δ45^ rat model, the CRISPR/Cas9 system was used to delete exon 45 of the rat dystrophin gene, which has an identical exon organization and reading frame to the human gene, including the mutational hotspot region spanning exons 45-55 (Ensembl Reference ENSG00000198947 and ENSRNOG00000046366, for the human and rat, respectively). A single-guide RNA (sgRNA) was designed to target exon 45 (**Figure 1A**), and SpCas9/gRNA vector was injected into Sprague Dawley zygotes. This approach exploited the error-prone non-homologous end joining (NHEJ) DNA repair mechanism to create a double-stranded break in exon 45. The resulting founder line carried a deletion of 606 bp encompassing exon 45, as confirmed by genotyping and Sanger sequencing (**Figure 1B–C**). Loss of dystrophin expression in skeletal and cardiac muscles from 3 weeks-old *Dmd*^Δ45^ rats, detected by western blot, confirmed disruption of the *Dmd* reading frame (**Figure 1D**). Immunostaining of extensor digitorum longus (EDL) skeletal muscle sections with dystrophin-specific antibodies targeting the C- and N-terminals further validated the absence of dystrophin expression (**Figure 1E**).

**Figure 1.**
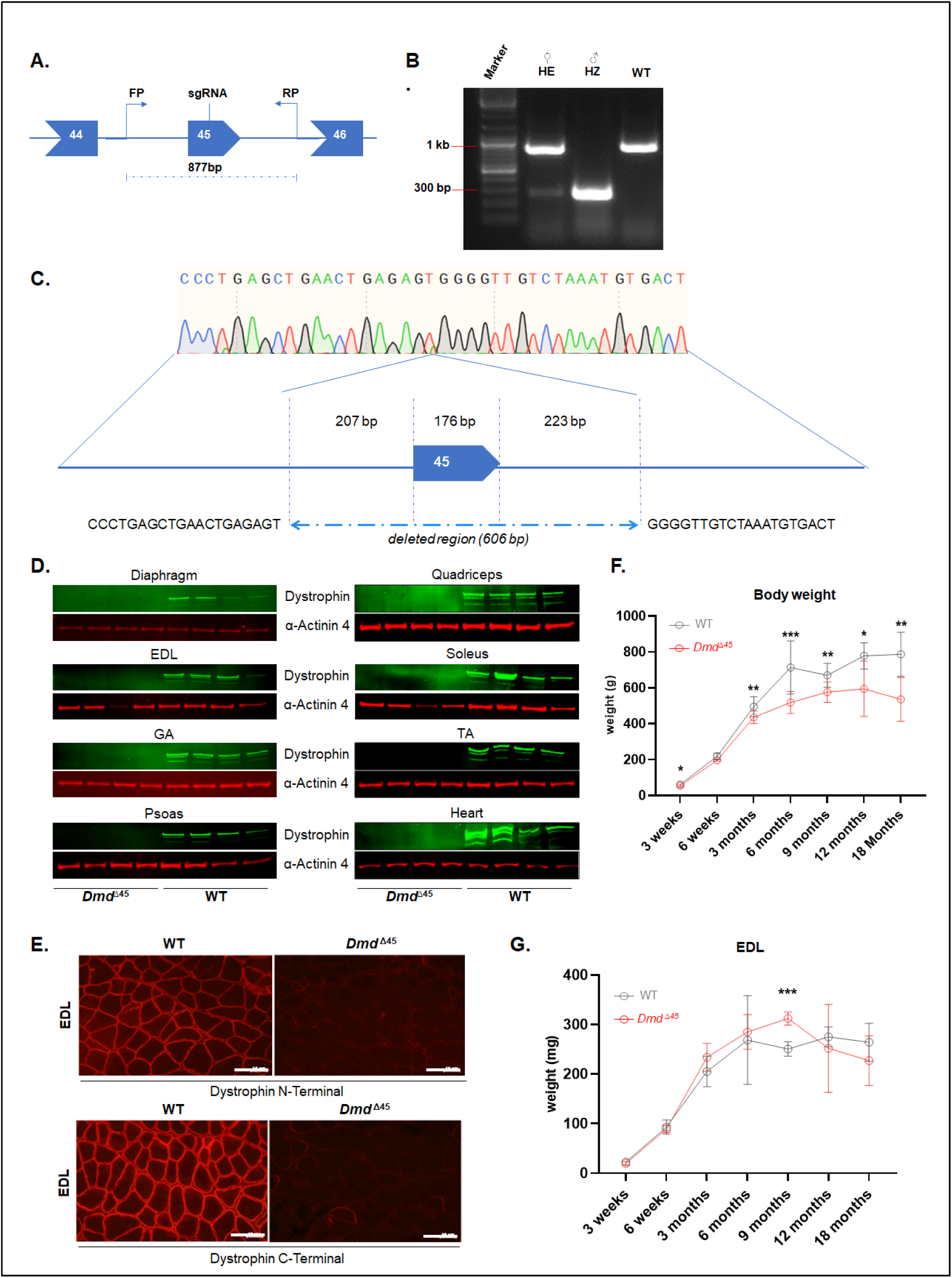
*Dmd^Δ45^* rats show absent dystrophin expression, progressive weight loss and late muscle atrophy. **(A)** Schematic representation of the targeted *Dmd* locus in *Dmd^Δ45^* rats using a CRISPR-based strategy with a single guide RNA. FP: forward primer, RP: reverse primer used for genotyping**. (B)** Genotyping of Dmd^Δ45^ rats. ♀HE female heterozygous; ♂ HZ male hemizygous, WT: wildtype. The apparent unequal amplification of methylated versus non-methylated DNA strands is likely due to X-inactivation in females as previously reported [47]. (**C**) Schematic representation of the 606 bp deletion in the *Dmd^Δ45^* rat model at the targeted locus. (**D**) Western blot analysis for dystrophin expression in Diaphragm, extensor digitorum longus (EDL), gastrocnemius (GA), psoas (Pso), quadriceps (Qua), soleus (Sol), tibialis anterior (TA), and heart tissues from 3-week-old Dmd^Δ45^ and WT rats (n=4 per group). **(E)** Representative images of EDL muscle cross-sections from 3-week-old WT and *Dmd^Δ45^* rats immunostained for dystrophin, using antibodies targeting the N- and C-terminal regions of dystrophin. Scale bar = 10 µM. **(F)** Body weight measurement of WT and *Dmd^Δ45^* rats from 3 weeks to 18 months (n= 3–16). **(G)** EDL muscle weight measurement of WT and *Dmd^Δ45^* rats from 3 weeks and 18 months (n=3-10).

The *Dmd*^Δ45^ rats exhibited moderately reduced lifespan. Among 146 wildtype (WT) and 151 *Dmd*^Δ45^ rats bred during the project, nine *Dmd*^Δ45^ rats died between 3 and 18 months of age, whereas no WT rats died before 18 months. Total body weights of *Dmd*^Δ45^ rats, recorded from 3 weeks to 18 months of age, were compared to age-matched WT controls (*n* = 3–16 per group). *Dmd*^Δ45^ rats displayed reduced weight gain starting at 3 weeks, which persisted throughout their lifespan (**Figure 1F**). Interestingly, analysis of individual skeletal muscle weights, including EDL, gastrocnemius (GA), tibialis anterior (TA) and soleus, showed no significant differences in weight between *Dmd*^Δ45^ rats and WT rats at early timepoints (3 weeks, 6 weeks, 3 months, and 6 months) (**Figure 1G and Figure S1**). Surprisingly, a significant increase in skeletal muscle weight was observed in *Dmd*^Δ45^ at 9 months, followed by a gradual decline between 9 and 18 months. In contrast, WT rats maintained stable skeletal muscle weights over this period. These findings suggest a late-onset, progressive muscle atrophy in the *Dmd*^Δ45^ rat model.

### Histological characterization shows key dystrophic features in the skeletal muscle

Next, we set out to characterize the histological features of skeletal muscle from early to later stages of disease progression. Hematoxylin Phloxine Saffron (HPS) and Sirius Red staining on quadriceps (QUA) muscles showed hallmark dystrophic features as early as 3 weeks of age. (**Figure 2A**). Notably, fibrosis was prominent from this early stage (**Figure 2A-B**), particularly in degenerative regions, suggesting a reactive fibrotic response to muscle breakdown. This was evidenced by light pink staining on Sirius Red, oedema-like regions on HPS staining, increased connective tissue deposition in HPS-stained sections (**Figure 2C**) and reduced muscle area (**Figure 2D**). Interestingly, fibrosis and connective tissue content showed a transient reduction at 3 months in quadriceps, potentially due to active muscle regeneration. However, progressive interstitial fibrosis developed thereafter, culminating in extensive fibrotic remodelling by 18 months, as observed in both Sirius Red and HPS staining, as well as in corresponding quantifications (**Figure 2A-C**). This fibrotic progression was also noted in other skeletal muscles, including the gastrocnemius (GA) and extensor digitorum longus (EDL) (**Figure S2A-B**). HPS staining additionally revealed substantial inflammatory infiltrates at 3 weeks (**Figure 2A, Figure S2A**). This peak in inflammation was partially resolved over time, not only in the quadriceps but also in other skeletal muscles (**Figure 2E, Figure S2B**). Immunostaining for CD68, a marker of monocyte lineage cells, confirmed these findings, with a high abundance of CD68+ infiltrates at 3 weeks, which diminished by 3 months and were scarce at 12 months (**Figure 2F**).

**Figure 2.**
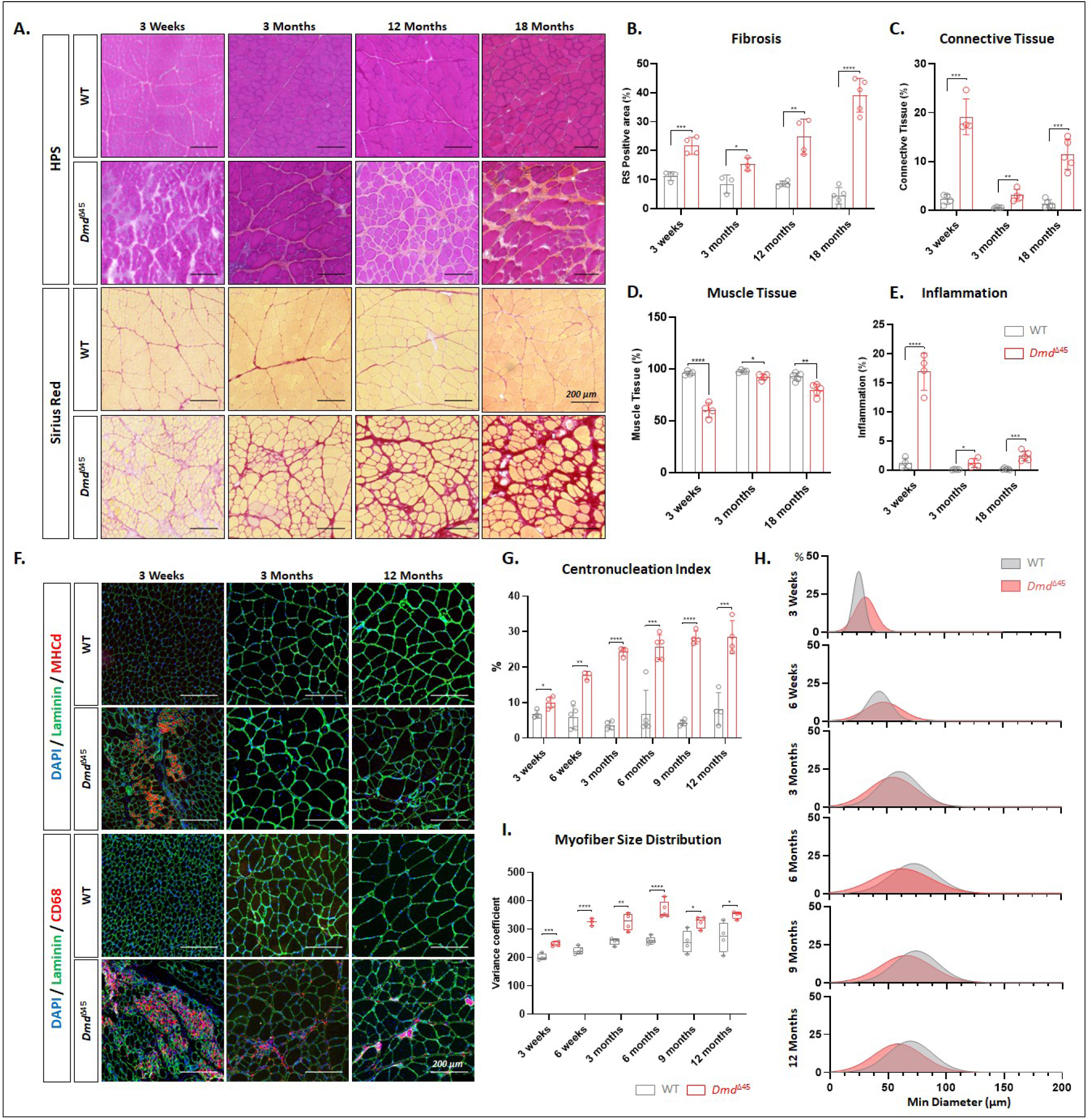
Histological evaluation of quadriceps muscles reveals key dystrophic features in *Dmd^Δ4^*^5^ rats. **(A)** Representative images of quadriceps cross-sections stained with Hematoxylin Phloxine Saffron (HPS) and Sirius Red (SR) from WT and *Dmd^Δ45^* rats at 3 weeks, 3 months, 12 months, and 18 months. **(B)** Quantification of fibrosis based on Sirius Red-positive area (n = 3– 5 rats per group, with 2–3 whole muscle cross-sections quantified per rat). **(C)** Quantification of connective tissue percentage from HPS-stained cross-sections (n = 3–5). **(D)** Quantification of muscle tissue percentage from HPS-stained cross-sections (n = 3–5). **(E)** Quantification of inflammatory areas in HPS images (n=3-5). **(F)** Representative images of quadriceps cross-sections immunostained for Laminin and developmental myosin heavy chain (MHCd) (upper panel) or Laminin and CD68 (lower panel). **(G)** Quantification of the centronucleation index based on Laminin and DAPI staining of cross-sections from rats between 3 weeks and 12 months (n = 3–5 per group, with 2–3 whole muscle cross-sections per rat). **(H)** Density plots depicting myofibre minimum diameter size distribution in WT and Dmd^Δ45^ rats between 3 weeks and 12 months. **(I)** Analysis of myofibre size distribution in the gastrocnemius (GA) muscle. Data are presented as box plots (median + minimum/maximum) showing the coefficient of variance, calculated as: [standard deviation of the muscle fibre size/mean of muscle fibre size] *1000.

To assess muscle regeneration, we performed immunostaining for developmental myosin heavy chain (MHCd) (**Figure 2F**). At 3 weeks, the presence of numerous MHCd+ myofibers suggested an early activation of muscle regeneration, likely triggered by the initial wave of muscle degeneration. This regenerative response diminished over time, as indicated by a marked reduction in MHCd+ myofibers. Concurrently, muscle tissue percentage increased between 3 weeks and 3 months (**Figure 2D, Figure S2B**). The centronucleation index, a key marker of muscle regeneration, increased with age and stabilized around 6 months in the quadriceps and other skeletal muscles (**Figure 2G, Figure S2C**), aligning with MHCd staining patterns that showed early positivity at 3 weeks but became sparse later, indicative of an initial robust regenerative phase that eventually plateaued.

Analysis of myofiber size revealed an early increase in *Dmd^Δ45^* rats at 3 weeks compared to WT (**Figure 2H**), followed by a progressive decline, ultimately leading to significantly smaller myofibers in dystrophic muscle. This pattern correlated with muscle weight analysis (**Figure 1G, Figure S1**) which showed an initial increase in muscle mass, followed by progressive loss, suggesting a late-onset muscle atrophy. Coefficient of variance analysis further highlighted increased heterogeneity in myofiber size distribution from 3 weeks onward (**Figure 2I**), a pattern consistent with disease progression.

Overall, the histological characterization of quadriceps and other skeletal muscles in *Dmd^Δ45^* rats revealed hallmark features of progressive muscular dystrophy. The disease course was characterized by an initial phase of muscle degeneration and inflammation, triggering an early regenerative response. However, this was followed by progressive muscle deterioration, marked by interstitial fibrosis and muscle atrophy.

### Elevated circulating biomarkers and decline in muscle function in *Dmd*^Δ45^ rats

To monitor disease progression, we assessed various serum biomarkers at multiple time points. Creatine kinase (CK), a widely used marker of myofiber damage, was measured first. CK levels (U/L) were significantly elevated in *Dmd*^Δ45^ rats by 3 months and increased further at 6 months before declining between 9 and 12 months (**Figure 3A**). This age-related decline in CK levels mirrors observations in patients, where CK decreases as motor function deteriorates and muscle mass is replaced by fibro-fatty tissue [3, 31]. We also measured circulating levels of Myomesin 3 (MYOM3) fragments, a more specific biomarker of myofibre damage in muscular dystrophies compared to CK [32]. MYOM3 levels were significantly elevated in *Dmd*^Δ45^ rats compared to WT controls starting at 3 weeks, with further increase observed up to 3 months (**Figure 3B**). Interestingly, MYOM3 levels showed a different trajectory from CK, decreasing at 6 months, and rising again at 9 and 12 months (**Figure 3A-B**).

**Figure 3.**
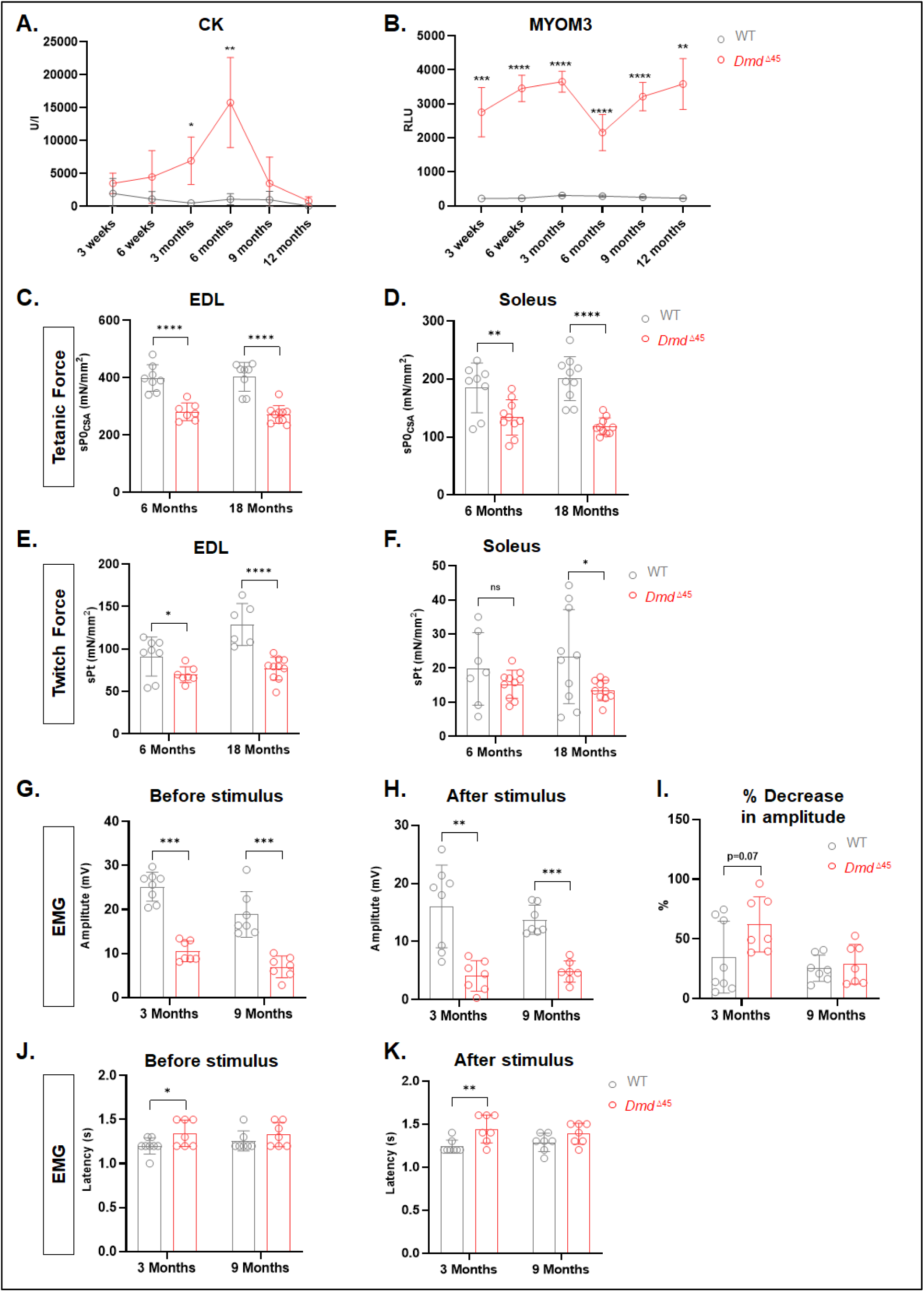
Elevated muscle damage biomarkers and impaired muscle force indicate muscle dysfunction in *Dmd^Δ45^* rats. **(A)** Measurement of serum creatine kinase (CK) activity in rats between 3 weeks and 12 months (n = 4–5 per group). **(B)** ELISA quantification of myomesin-3 (MYOM3) fragments in rat serum between 3 weeks and 12 months (n = 4–5 per group). **(C–F)** In vitro mechanical force measurements in EDL and soleus muscles from 6- and 18-month-old rats (n = 7–10 per group). sP0csa: maximal tetanic force normalized to muscle cross-sectional area; sPt: maximal twitch force normalized to muscle cross-sectional area. **(G–H)** Electromyography (EMG) analysis of the GA muscle in 3- and 9-month-old rats, showing the amplitude of the compound muscle action potential (CMAP) before and after stimulus trains (4 × 200 stimuli at 10 Hz) (n = 7–8 per group). **(I)** Bar plot depicting the percentage decrease in CMAP amplitude following stimulus. **(J–K)** Bar plots showing latency, defined as the time required for the electrical impulse to travel from the nerve to the muscle, before and after stimulus.

To evaluate skeletal muscle function in the *Dmd*^Δ45^ rat model, mechanical force generation was measured *ex vivo* in EDL and soleus muscles isolated from 6- and 18-month-old *Dmd*^Δ45^ rats. Tetanic force, which represents the maximum sustained force generated by a muscle during continuous, high-frequency electrical stimulation, was measured first. In the EDL, *Dmd*^Δ45^ rats exhibited a significant reduction in tetanic force compared to WT controls at both ages, generating approximately 70% of the average maximum tetanic force normalized to muscle cross-sectional area produced by WT muscle (**Figure 3C**). In the soleus, however, the reduction was less pronounced at 6 months (134 mN·mm⁻² in *Dmd*^Δ45^ vs. 185 mN·mm⁻² in WT) but became more pronounced by 18 months (124 mN·mm⁻² in *Dmd*^Δ45^ vs. 201 mN·mm⁻² in WT) (**Figure 3D**). Twitch force, defined as the force generated by a single, brief contraction in response to a single electrical stimulus, was also assessed. In EDL muscles, *Dmd*^Δ45^ rats showed significantly reduced maximum twitch force at both 6 months (70 mN·mm⁻² in *Dmd*^Δ45^ vs. 91 mN·mm⁻² in WT) and 18 months (77 mN·mm⁻² in *Dmd*^Δ45^ vs. 129 mN·mm⁻² in WT) (**Figure 3E**). In the soleus, there was no significant difference in twitch force between the groups at 6 months (**Figure 3F**). However, by 18 months, a significant reduction in twitch force was observed in *Dmd*^Δ45^ rats (13 mN·mm⁻² in *Dmd*^Δ45^ vs. 23 mN·mm⁻² in WT) (**Figure 3F**). It is noteworthy that the soleus muscle contains a majority of slow-twitch myofibers, unlike the EDL, which predominantly consists of fast-twitch fibres. This composition likely explains why the soleus muscle is less affected at 6 months. Overall, these findings reveal a progressive decline in both tetanic and twitch force in *Dmd*^Δ45^ rats, suggesting a gradual loss of the muscle’s capacity for sustained force production during prolonged contractions and its ability to respond to rapid, isolated activations.

Furthermore, to gain insights into neuromuscular activity, we performed electromyography (EMG) on the GA muscle to evaluate its response to sciatic nerve stimulation. The amplitude of the motor response, recorded as the compound muscle action potential (CMAP), reflects the number of activated muscle fibres. CMAP amplitude was measured both before and after stimulus trains (4 × 200 stimuli at 10 Hz). In *Dmd*^Δ45^ rats, the CMAP amplitude, both before and after the stimulus trains, was significantly reduced compared to WT rats at both 3 and 9 months of age (**Figure 3G-H).** At 3 months, *Dmd*^Δ45^ rats exhibited a greater percentage decrease in CMAP amplitude following stimulus trains, indicative of increased muscle fatigue (**Figure 3I**). However, this difference did not reach statistical significance (p = 0.0721). By 9 months, the percentage decrease in CMAP amplitude after stimulation was comparable between *Dmd*^Δ45^ and WT rats (**Figure 3I**). Latency, which measures the time required for the electrical impulse to travel from the nerve to the muscle and provides insights into nerve conduction, was significantly increased in *Dmd*^Δ45^ rats at 3 months (both before and after stimulus trains) (**Figure 3J-K**). However, no significant difference in latency was observed between genotypes at 9 months (**Figure 3J-K**). These findings reveal a reduction in neuromuscular activity, responsiveness to stimulation, and evidence of neuropathy in *Dmd*^Δ45^ rats. Interestingly, the observed defects were more pronounced at 3 months compared to 9 months, suggesting a potential adaptive response in the neuromuscular system over time.

### Cardiac histological and functional assessments

We next evaluated heart structure and function in *Dmd^Δ45^* and WT control rats at various ages to characterize the cardiac pathology, which is currently recognized as the leading cause of death in DMD patients [13, 33]. Typically, DMD patients develop a progressive dilated cardiomyopathy, marked by inflammatory cell infiltration, necrosis, excessive cardiac fibrosis [34], left ventricular dilation, and progressive decline in cardiac function, ultimately leading to heart failure. To investigate this phenotype, we first monitored heart tissue weights throughout the rat lifespan. Starting at 6 months, *Dmd^Δ45^* rats exhibited a significant increase in heart weight normalized to body weight, which persisted at 9 and 12 months and reached a 1.6-fold increase compared to WT controls by 18 months (**Figure 4A**).

**Figure 4.**
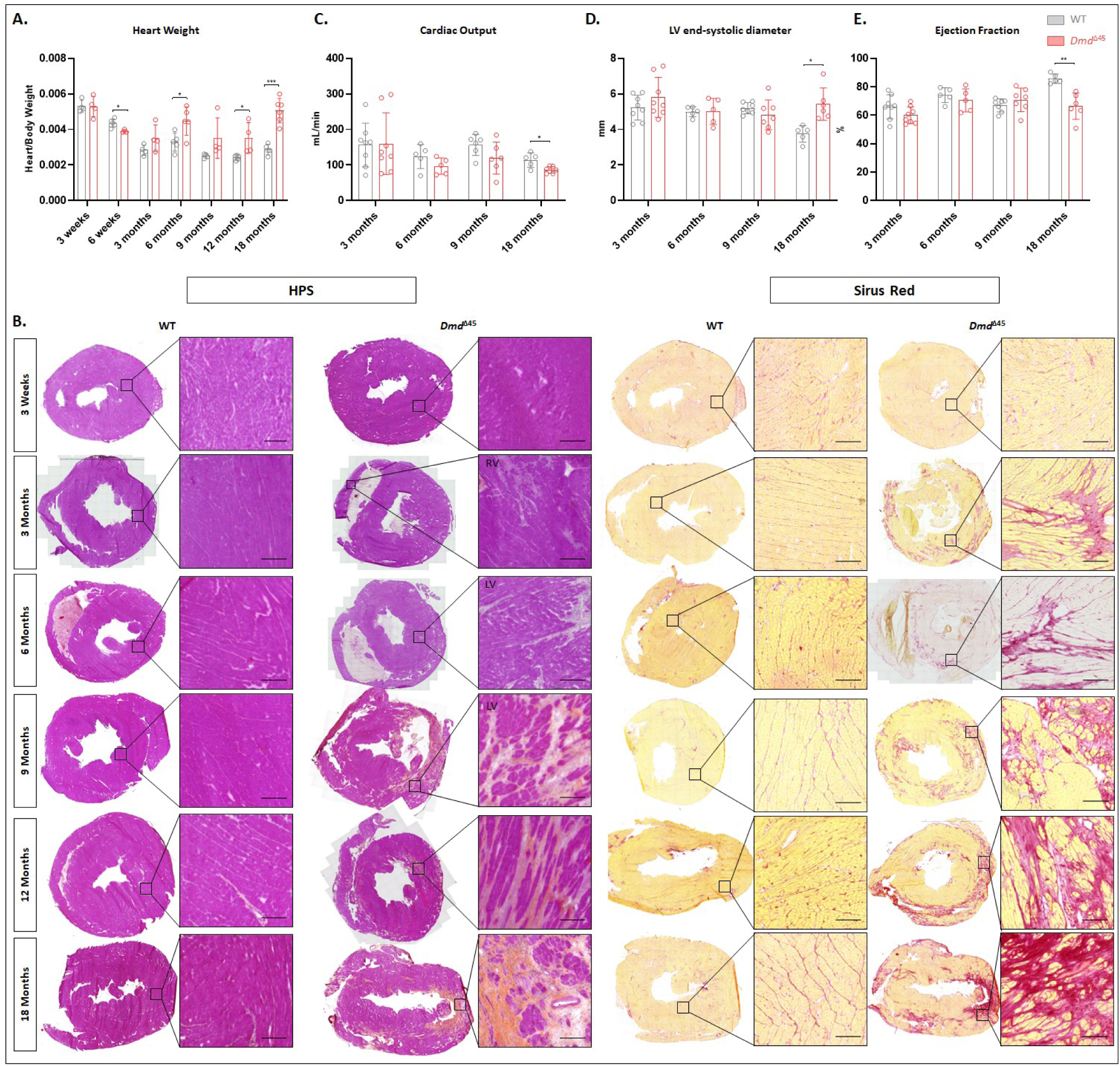
Cardiac characterization reveals progressive cardiac pathology in *Dmd^Δ45^* rats. **(A)** Heart weight normalized to body weight in rats from 3 weeks to 18 months (n = 3–6 per group). **(B)** Histological characterization of heart tissues from WT and *Dmd^Δ45^* rats between 3 weeks and 18 months. The left panel shows HPS staining, while the right panel displays Sirius Red staining highlighting cardiac fibrosis (Bar plot = 200µm). **(C)** Cardiac output measured by transthoracic echocardiography (TTE) in WT and *Dmd^Δ45^* rats between 3 and 18 months of age (n=5-8). **(D)** Left ventricular (LV) end-systolic diameter measurements obtained via TTE (n=5-8). **(E)** Bar plot showing ejection fraction quantification by TTE in rats from 3 to 18 months of age (n=5-8).

Heart histology was assessed from 3 weeks to 18 months using HPS and Sirius Red staining to evaluate tissue remodelling and abnormalities (**Figure 4B**). At 3 weeks, no notable abnormalities were observed. However, by 3 months, inflammatory infiltrates and reactive interstitial fibrosis were evident, predominantly in the epicardial region of the right ventricle (RV). Between 6 and 9 months, significant structural changes emerged in the left ventricle (LV) and septum, including pronounced inflammation and dense replacement fibrosis, with the LV being more severely affected than the RV. Fibrosis deposits progressively increased until 18 months, at which point a substantial portion of the heart tissue, including both the RV and LV, was composed of dense fibrotic networks.

Cardiac function and structure were further evaluated through transthoracic echocardiography at 3, 6, 9, and 18 months, with colour Doppler imaging performed at 3 and 9 months to gain additional insights into cardiac flow patterns, velocities, and pressures (**Table S1**). At 3 months of age, no significant changes in LV structure or function were observed in *Dmd^Δ45^* rats. However, a significant increase in pulmonary artery (PA) and pulmonary valve (PV) parameters was noted, including increased PA velocity, pressure gradient, pressure, and pulmonary ejection time compared to WT controls. These findings indicate pulmonary hypertension, predictive of right heart failure. This correlates well with histological observations showing greater pathological changes in the RV than the LV at this age (**Figure 4B**). At 6 months of age, no significant changes in systolic function were observed, but there was a slight (albeit non-significant) reduction in stroke volume and cardiac output, possibly due to reduced RV preload (**Figure 4C**). At 9 months of age, echocardiography and Doppler imaging revealed a decrease in several LV hemodynamic parameters, including aortic velocity, pressure gradient, pressure, stroke volume, and cardiac output. These changes were indicative of LV diastolic dysfunction, characterized by prolonged LV relaxation time (IVRT) and increased LV filling pressures (**Table S1**). These abnormalities led to tricuspid valve insufficiency, RV failure, and signs of heart failure with preserved ejection fraction (HFpEF). At 18 months of age, significant decreases in ejection fraction (EF) and increased LV end-systolic diameter (LVESD) indicated systolic dysfunction with decreased contractility (**Figure 4D-E**), consistent with heart failure with reduced ejection fraction (HFrEF). Increased end-systolic volume (ESV) and significantly reduced cardiac output were observed (**Figure 4C**). However, LV end-diastolic diameter (LVEDD) and end-diastolic volume (EDV) did not increase, indicating the absence of compensatory mechanisms, such as increased filling. Overall, the cardiac characterization of *Dmd^Δ45^*rats reveal a progressive cardiac pathology that mirrors key features observed in DMD patients, including the transition from RV-dominant abnormalities and pulmonary hypertension in early stages to LV dysfunction and heart failure at later stages.

### Behavioural evaluation of *Dmd*^Δ45^ rats show cognitive impairments

Cognitive impairments, including non-progressive deficits with diverse manifestations, have been reported in many DMD patients [35–37]. To evaluate behavioural and cognitive function, as well as locomotor activity, in *Dmd*^Δ45^ rats, we conducted a series of tests on 3-month-old rats. The open field test, a widely used assay for assessing exploratory behaviour, general locomotor activity, and anxiety-related responses, revealed significant differences between *Dmd*^Δ45^ rats and WT controls. *Dmd*^Δ45^ rats exhibited a significantly decreased total distance travelled (**Figure 5A**), indicating reduced exploratory behaviour and locomotor activity. Additionally, they spent less time in the centre of the arena and more time along the periphery (**Figure 5B**), suggesting increased anxiety or decreased curiosity. However, no differences were observed in the number of rearing events (**Figure 5C**), indicating some aspects of exploratory behaviour remained intact.

**Figure 5.**
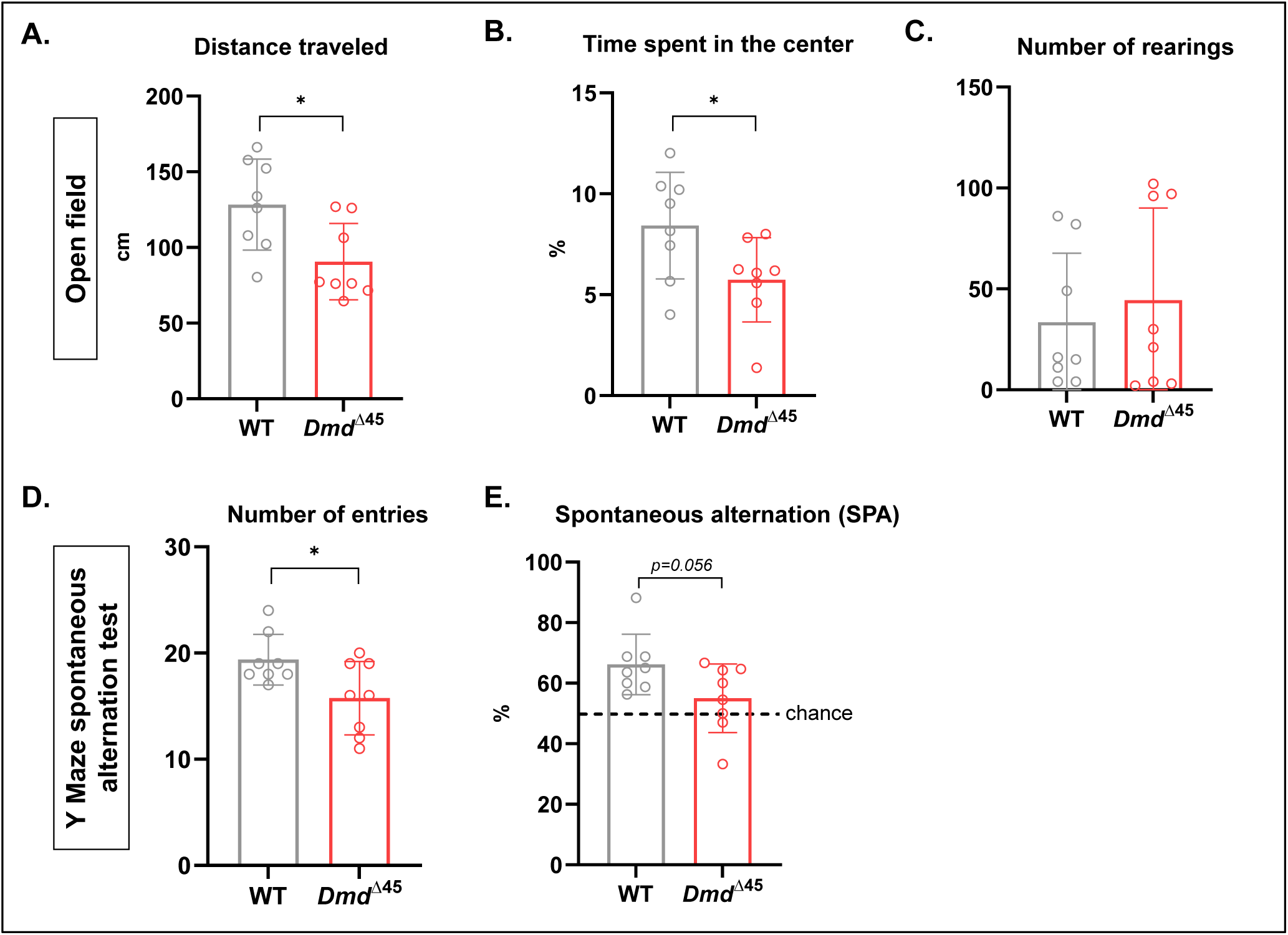
Behavioural assessment reveals cognitive impairments in *Dmd^Δ45^* rats. (A-C) Open field test in 3-month-old WT and *Dmd^Δ45^* rats, analysing total distance travelled **(A)**, percentage of time spent in the centre **(B)**, and number of rearings **(C)** (n = 8 per group). **(D-E)** Y-maze spontaneous alternation test, quantifying the total number of arm entries **(D)** and the spontaneous alternation (SPA) percentage between arms **(E)**.

To assess spatial and working memory, we employed the Y-maze spontaneous alternation test, in which rats were allowed to explore a Y-shaped maze freely for 8 minutes. *Dmd*^Δ45^ rats showed a reduced number of arm entries, consistent with decreased locomotor activity and signs of anxiety (**Figure 5D**). Furthermore, their spontaneous alternation percentage—a measure of working memory based on the proportion of consecutive entries into all three arms without repetition—was lower than WT controls (**Figure 5E**). Although this decrease did not reach statistical significance (p = 0.056), the alternation percentage was significantly higher than chance only (50%) in WT rats, indicating impaired working memory. Overall, these findings suggest that *Dmd*^Δ45^ rats exhibit cognitive and behavioural impairments, including reduced exploratory behaviour, heightened anxiety-related responses, and deficits in working memory, which align with previously reported symptoms in other DMD rat models [29].

### RNA sequencing analysis indicates dysregulated pathways related to *DMD*

To gain insight into the consequences of exon 45 deletion in *Dmd*^Δ45^ rats at molecular level, we performed RNA sequencing (RNAseq) on skeletal (Pso, Sol and Dia) and cardiac muscles from both 6- and 12-month-old rats (complete data are provided in the **Supplemental Excel file**). First, we quantified the reads of each exon of the *Dmd* gene, confirming the deletion of exon 45 (**Figure S3A**) and examined the expression of the *Dmd* gene, which showed a dramatic decrease in all muscle samples (**Figure S3B**). We further analysed exon usage in the *Dmd* gene by grouping exons into two regions: “Beginning” (upstream of exon 45) and “End” (downstream of exon 45). Consistent with previously observed transcriptional imbalance in DMD patients [38], we found a progressive reduction in *Dmd* transcript levels decreased from the Beginning to the End exons (**Figure S3C**). Given that utrophin (*Utrn*) can partially compensate for the loss of dystrophin, we assessed its expression and observed significant upregulation over time in skeletal muscles, with a modest, non-significant increase in the heart (**Figure S3D)**.

We then performed principal component analysis (PCA) on skeletal muscle tissues to reduce data complexity. PCA revealed a clear separation between *Dmd*^Δ45^ and WT groups, with PC1 capturing genotype-driven differences and PC2 reflecting both genotype and tissue-specific variation (**Figure 6A**). Notably, the transcriptomic divergence *Dmd*^Δ45^ and WT samples exceeded the variation among different skeletal muscle types. To identify genes contributing most to these differences, we averaged the PC1 contribution scores (i.e. the weights indicating how strongly each gene influences PC1) across all skeletal muscle RNA-seq datasests at both timepoints. Genes with an average contribution score >0.03 were selected, representing those most strongly associated with genotype-driven transcriptional changes across muscles and timepoints. This approach identified key 36 genes (**Figure 6B**, **Table S2). Approximately,** half of these had been previously reported as dysregulated in DMD such as *MymX*, *Mymk*, *Spp1*, *Thbs4*, *Postn*, *Myl4*, *Timp1* and *Aqp4*, while the others were novel in this context (e.g *Igfn1, Draxin, Lrrc15, TMEM184a, Ptx4 and Cilp*, for the upregulated genes and *Barx2* and *Mylk4* for the down regulated ones). The dot-graph of gene expression for these 36 genes in skeletal muscles illustrated that 33 genes were up-regulated and 3 genes were down-regulated in *Dmd*^Δ45^ rats compared to WT rats (**Figure 6B**). Of note, applying the same selection criteria to PC2 genes revealed that all top PC2 contributors were already included in the PC1-derived 36-gene list. A Volcano plots highlighting globally dysregulated genes (fold-change> 2 ; adjusted p-value < 0.05) are shown in **Figure S4A.** Among skeletal muscles, the diaphragm exhibited the most pronounced dysregulation (with FC exceeding 1000), followed by the psoas and soleus, in agreement with histological findings. Venn diagrams summarize the overlap of differentially expressed genes (DEGs) across muscles at both time points (**Figure 6C)**. At 6 months, 179 were upregulated genes across three muscles, with 56 additional genes shared by psoas and soleus. By 12 months, shared upregulated genes rose to 625, with 797 more shared between psoas and soleus. For downregulated genes, 25 genes were common across all muscles at 6 months (excluding *Dmd*), with 7 genes shared by psoas and soleus. By 12 months, this number increased to 58 common downregulated genes, and 34 shared between psoas and soleus. A temporal analysis of DEG evolution (**Figure 6D**, **Tables S3** and **Figure S4A)** suggested an overall age-dependent worsening, with the exception of the diaphragm.

**Figure 6:**
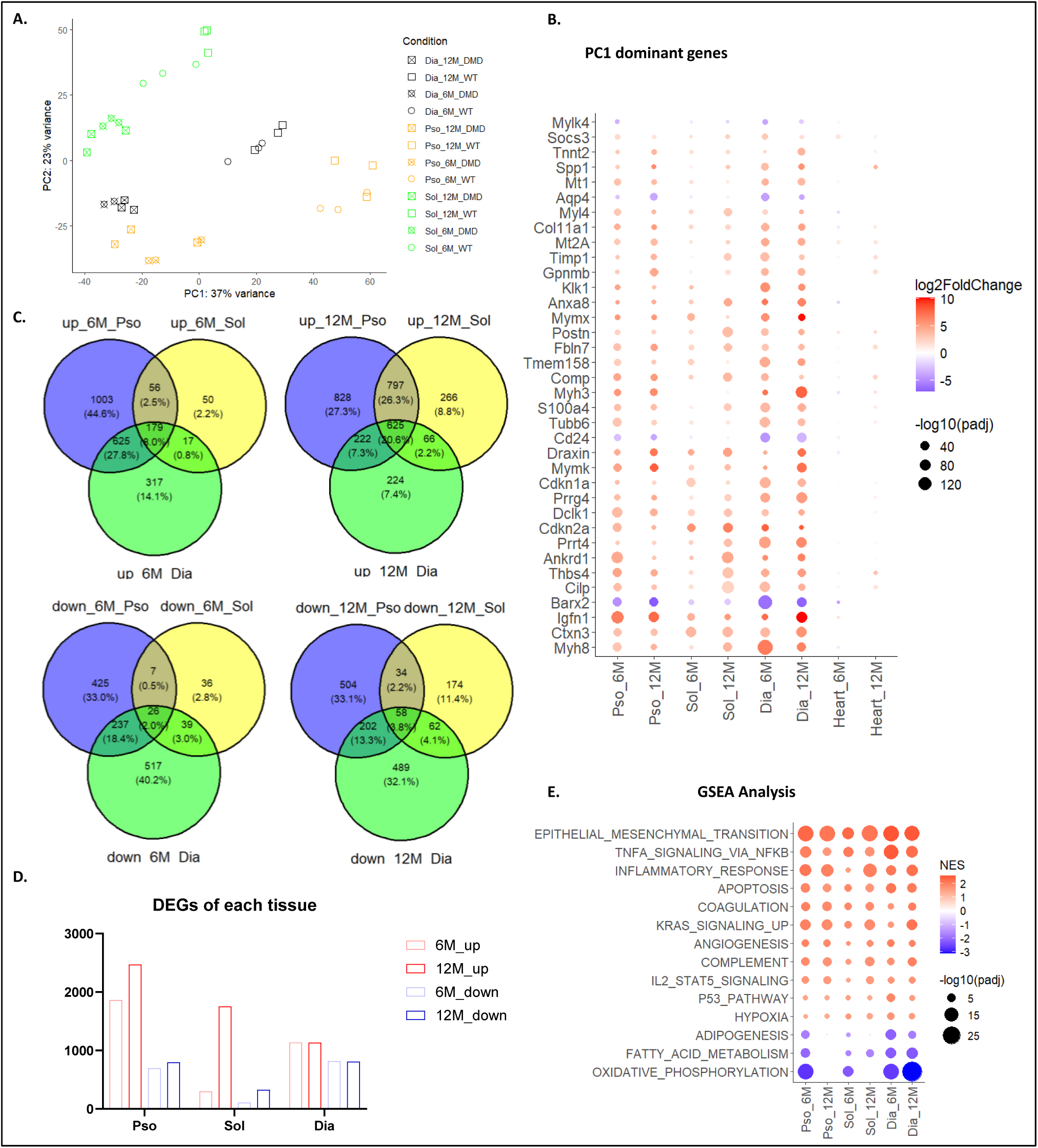
RNAseq analysis revealed the dysregulated pathways related to DMD. (**A**) Principal Component Analysis (PCA) scatter plot of gene expression profiles. Dia samples are shown in black, Pso in orange, Sol in green. Samples collected at 6M are presented by circles, and those at 12M by squares. DMD rats are indicated with a cross overlaid on the WT sign. (**B**) Dot graph of the PC1 dominant genes. Each dot represents a gene in a specific *Dmd^Δ45^* sample. Colour indicates the log₂ fold change, with blue for downregulation and red for upregulation. Dot size reflects the -log10(padj) for that gene in the corresponding *Dmd^Δ45^*sample. Genes are ordered based on the geometric mean of their adjusted p-values across all conditions, highlighting the most consistently significant genes. The result showed that the heart does not present similar dysregulation than in skeletal muscles. (**C**) Venn diagrams illustrating the overlap of commonly modified genes across different conditions. The comparison between WT and *Dmd^Δ45^* is shown in blue for the psoas, yellow for the soleus, and green for the diaphragm. (**D**) Bar plot illustrating the number of the Differentially Expressed Genes (DEGs) among skeletal muscles. (**E**) Dot graph of the commonly enriched pathways. Each dot represents a pathway in a specific *Dmd^Δ45^* sample. Dot size indicates the log_10_(padj). Dot colour reflects the Normalized Enrichment Score (NES) for that pathway in the corresponding *Dmd^Δ45^* sample. Pathways are ordered based on the average of their NES across all conditions.

We performed Gene Set Enrichment Analysis (GSEA) on skeletal muscle RNA-seq using Hallmark gene sets from the MSigDB database (adjusted p < 0.05) (**Figure 6E, Figure S4B**). Overall, the most significantly dysregulated hallmark pathways were upregulated and included epithelial to mesenchymal transition, inflammation and immune responses, and stress- or death-associated processes (e.g., apoptosis, DNA repair) (**Figure 6E**). In contrast, downregulated pathways primarily involved metabolic processes, such as adipogenesis, fatty acid metabolism, and oxidative phosphorylation (**Figure 6E**). At 12M, the upregulated pathways remained largely consistent, while downregulation became more concentrated on fatty acid metabolism.

Regarding the heart, PCA analysis revealed no clear separation between WT and *Dmd*^Δ45^ rats at 6 months. However, at 12 months, the transcriptomic profiles of two out of three *Dmd*^Δ45^ rats were distinct from WT controls, potentially reflecting inter-individual variability in the onset or progression of cardiomyopathy (**Figure 7A)**. Differential expression analysis identified 35 and 231 upregulated genes, and 24 and 76 downregulated genes in *Dmd*^Δ45^ hearts at 6 and 12 months, respectively, compared to age-matched WT controls (**Figure 7B, Table S5**). The DEGs were enriched in pathways related to inflammation and immune responses (e.g., *Cxcl1*, *Socs3*, *Nfkbiz*, *Selplg*), fibrosis and ECM remodelling (*Thbs1*, *Adamts1*, *Serpine1*, *Ccn1*) and transcriptional regulation (*Nr4a1/2*, *Bhlhe40*). Conversely, several downregulated genes were involved in muscle structure and function (*Neb*) and metabolism (*Adh6*, *Pxmp4*).

**Figure 7:**
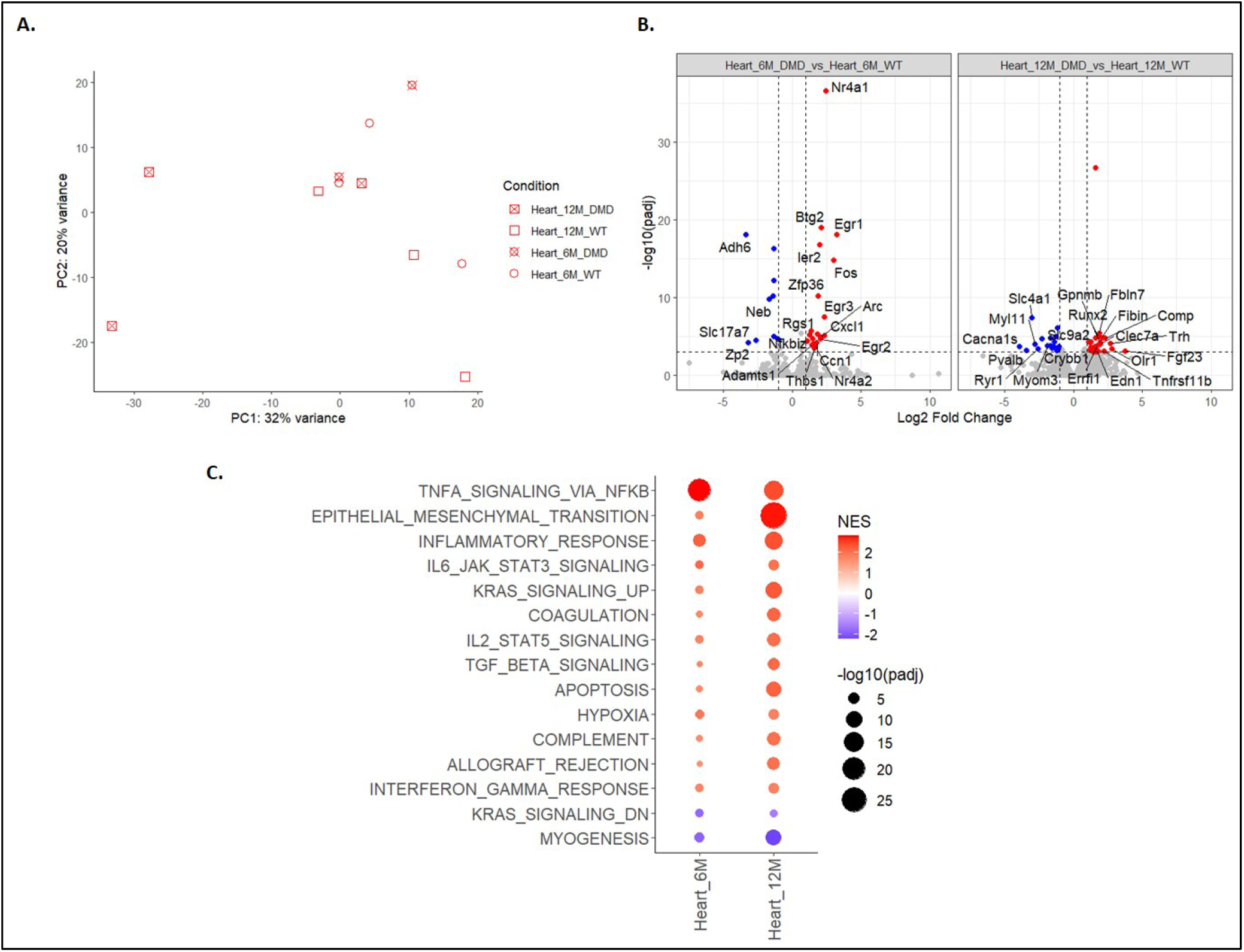
RNA-seq analysis revealed the dysregulated pathways in heart. (**A**) PCA scatter plot of gene expression profiles. Samples collected at 6M are presented by circles, and those at 12M by squares. *Dmd^Δ45^* rats are indicated with a cross overlaid on the WT sign. (**B**) Volcano plots showing differential gene expression in heart samples, comparing DMD and WT rats at 6M and 12M. (**C**) GSEA highlighting differentially enriched Hallmark pathways in heart samples, comparing *Dmd^Δ45^*and WT rats at 6M and 12M.

Interestingly, three genes upregulated in the heart at 12 months, *Comp*, *Gpnmb*, and *Fbln7*, were also part of the PC1 gene set previously identified in skeletal muscle. *Comp* has been reported as a biomarker of fibrosis in a DMD rat model [27], while *Gpnmb* is both a marker and effector of growth factor-expressing macrophages and contributes to muscle regeneration in DMD [39]. In contrast, *Fbln7*, which encodes Fibulin-7, a member of the ECM-associated fibulin family [40], has not been previously associated with DMD. Notably, *Fbln7* was also upregulated in all skeletal muscles at both 6 and 12 months, suggesting its potential as a novel biomarker of ECM remodelling in DMD.

GSEA analysis further revealed dysregulated pathways in the heart (**Figure 7C**). At 6 months, there was significant activation of immune-related pathways, with TNFα signalling and inflammatory response among the most enriched. In contrast, pathways associated with myogenesis were downregulated. By 12 months, immune activation persisted but was less pronounced compared to 6 months, while pathways related to ECM remodelling became more prominent. These included epithelial-mesenchymal transition, apoptosis, and TGF-β signalling. Notably, the pathways altered in the heart at 12 months closely resembled those observed in the Psoas muscle at 6 months, suggesting that cardiac disease progression lags behind skeletal muscle pathology.

### Spontaneous exon skipping partially restore dystrophin expression in *Dmd^Δ45^* rat model

Although *Dmd*^Δ45^ rats exhibit skeletal muscle dystrophy and cardiomyopathy, molecular, histological, and functional analyses suggested a milder phenotype compared to other published rat models such as the exon 23 deleted rat [29] or *Dmd*^Δ52^ rat model [27]. Since all these models involve out-of-frame mutations in the *Dmd* gene, , we sought to understand the basis for the phenotypic differences observed. Notably, we detected rare sporadic dystrophin-positive fibres in EDL muscle sections from 3-week-old *Dmd*^Δ45^ rats. This prompted an investigation of revertant fibres across ages, from 3 weeks to 12 months. Quantification of revertant fibres revealed an age-dependent increase in dystrophin-positive fibres in skeletal muscles (**Figure 8A-B**, **Figure S5A-B**). In quadriceps, the percentage of revertant fibres reached up to 36% by 9 months of age (**Figure 8B**). In TA and soleus muscles, revertant fibres were also observed, though to a lesser extent —reaching up to 19.6% (± 6.5%) and 14.6% (± 6.5%), respectively, at 9 months. Western blot analysis further confirmed the re-emergence of dystrophin expression in skeletal muscle but not in the heart of *Dmd*^Δ45^ rats (**Figure 8C**).

**Figure 8:**
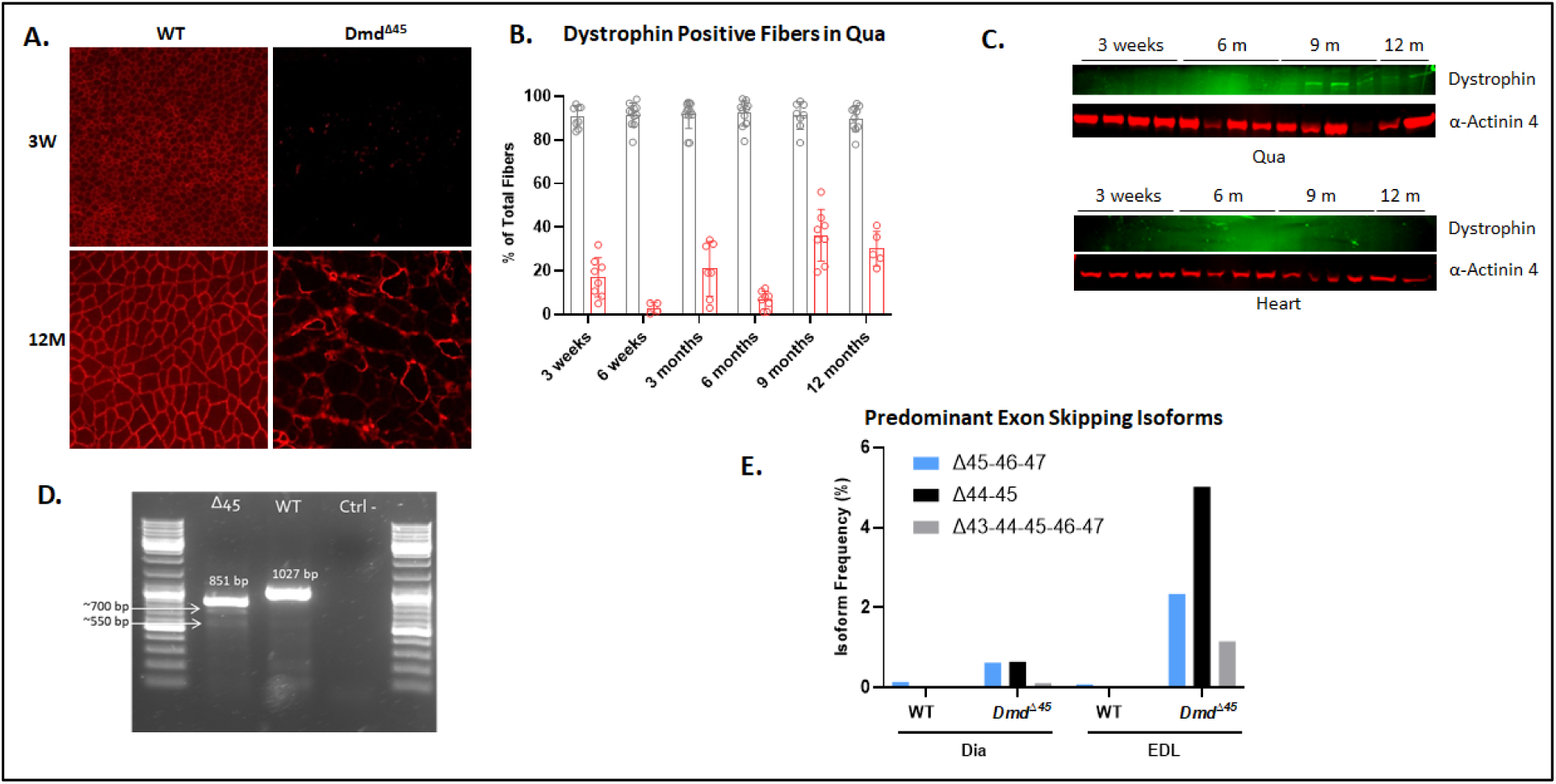
Spontaneous exon skipping partially restore dystrophin expression in Dmd^Δ45^ rat model. (**A**) Immunostaining for dystrophin in EDL Size bar=100 µM. (**B**) Quantification of dystrophin positive myofibres in different muscles from 3 weeks to 12 months of age in quadriceps muscle. (**C**) Western blot for Dystrophin in quadriceps and heart. m= month. (**D**) Proportion of observed exon skipping events.

Spontaneous exon skipping, which underlines the formation of revertant fibres, has been reported in patients with deletion of exon 45 in the *DMD* gene [41–43]. To investigate this mechanism in our model, we performed RT-PCR spanning exons 42 to 48 on skeletal muscle cDNA from 6-months-old WT and *Dmd*^Δ45^ rats. In WT samples, a single 1027 bp band corresponding to the full-length sequence between exons 42 and 48 was observed (**Figure 8D**). In *Dmd*^Δ45^ rat, in addition to the expected 851 bp band lacking exon 45, additional lower molecular weights bands (∼550 and 700bp) were detected (**Figure 8D**). **Sanger sequencing of these revealed** multiple exons-skipping events across muscles, many of which restored the open reading frame (**Figure S5C**). To confirm and quantify these events, we employed PacBio® long-read sequencing on exon 42-48 amplicons from WT and *Dmd*^Δ45^ EDL. In *Dmd*Δ45 rats, the major isoform involved skipping of exon 44 (5.02% of transcripts), followed by skipping of exons 46-47 (2.34%) and a larger multi-exon skip from exon 43 to 47 (1.16%) included (**Figure 8E**). Altogether, these findings demonstrate that the *Dmd*^Δ45^ model recapitulates spontaneous exon-skipping events similar to those reported in patients with exon 45 deletions

To explore the molecular mechanism underlying exon skipping in skeletal muscles of *Dmd*^Δ45^ rats, we analysed the expression of spliceosome-related genes from our RNA-seq dataset using the MSigDB gene set. We dientifed 49 splicesome-related gens that genes were significantly dysregulated (padj < 0.05), the majority of which were downregulated in skeletal muscle of *Dmd^Δ45^* rats (**Figure S5D**). In contrast, none of these genes were differentially expressed in the heart at either 6 or 12 months —consistent with the absence of revertant fibers in cardiac tissue. This suggests that a general downregulation of spliceosome components may lead to impaired spliceosome assembly, facilitating exon skipping selectively in skeletal muscle of *Dmd^Δ45^*rats.

## DISCUSSION

In this study, we generated a *Dmd*^Δ45^ rat model by deleting exon 45 of the *Dmd* gene, which represents the most frequent single-exon deletion within the mutational hotspot region in DMD patients. This deletion results in an out-of-frame mutation, leading to absence of dystrophin expression in both skeletal and cardiac muscle tissues by 3 weeks of age.

Phenotypically, *Dmd*^Δ45^ rats exhibited a significant increase in skeletal muscle mass at early time points, followed by a progressive decline between 9 and 18 months of age, suggesting the presence of an initial pseudohypertrophy followed by a late-onset progressive muscle atrophy. Histopathological analysis revealed hallmark features of muscular dystrophy, characterized initially by muscle degeneration and inflammation, which triggered a robust regenerative response. However, this was gradually replaced by progressive muscle deterioration, including interstitial fibrosis and marked muscle atrophy.

This structural deterioration was also reflected functionally. *Dmd*^Δ45^ rats demonstrated a progressive decline in both tetanic and twitch force, indicating a reduction in the muscle’s capacity for sustained force production as well as its responsiveness to brief activations. In parallel, we observed a decrease in neuromuscular activity and responsiveness to stimulation, along with evidence of neuropathy. Interestingly, these neuromuscular deficits were more pronounced at 3 months than at 9 months, suggesting a potential adaptive or compensatory response within the neuromuscular system over time.

Circulating biomarkers of muscle damage, including CK and MYOM3, were significantly elevated in *Dmd^Δ45^* rats compared to wild-type controls. Notably, these markers followed distinct temporal trajectories. CK levels peaked around 6 months and then gradually declined, whereas MYOM3 levels decreased at 6 months before increasing again thereafter. This divergence may reflect their underlying biological roles: CK, a cytosolic enzyme, is released upon sarcolemmal damage and mirrors early acute injury, while MYOM3, a structural protein, may better indicate muscle fiber necrosis and chronic damage progression.

Cardiac assessment revealed a progressive cardiomyopathy that recapitulates key clinical features of DMD, including a shift from early right ventricular (RV) dysfunction and pulmonary hypertension to later-stage left ventricular (LV) failure, accompanied by progressive fibrosis and cardiac remodelling. These cardiac manifestations are absent in *mdx* mice but have been consistently observed in all previously described DMD rat models. Beyond muscular and cardiac phenotypes, *Dmd^Δ45^* rats also exhibited cognitive and behavioural deficits, including reduced exploratory behaviour, increased anxiety-like responses, and impairments in working memory. These neuro-behavioural alterations parallel the cognitive symptoms observed in DMD patients. Overall, this model recapitulates main aspects of the clinical symptoms observed in DMD patients, confirming the rat suitability as a preclinical model mirroring the human disease more than existing mouse models.

Compared to previously reported DMD rat models, *Dmd^Δ45^*exhibited phenotypes that were similar to or more severe than those observed *Dmd*^Δ3-16^ and *Dmd*^mdx^ rats, but less severe than in *Dmd*^Δ52^ rats, which are characterized by rapid disease progression and early mortality [27]. Specifically, the reduction in weight gain occurred earlier in the *Dmd^Δ45^* rats than in *Dmd*^Δ3-16^ and *Dmd*^mdx^ rats but at a similar time point as in *Dmd*^Δ52^ rats [27–29]. In terms of circulating biomarkers, CK levels in *Dmd^Δ45^* rats were intermediate between those observed in *Dmd*^Δ3-16^/*Dmd*^mdx^ and *Dmd*^Δ52^ rats [27–29], further supporting the notion of an intermediate disease severity. Cardiac abnormalities in *Dmd*^Δ52^ rats were comparable in magnitude to those reported in *Dmd^mdx^* and *Dmd*^Δ3-16^ rats [44], suggesting that the *Dmd^Δ45^* model retains the ability to develop the full spectrum of DMD-associated cardiac pathology, which is often absent in mice models.

The intermediate phenotype presented by the *Dmd^Δ45^* may be attributed to the presence of a relatively high number of revertant fibres, reaching up to 36% in the quadriceps by 9 months of age. This contrasts with *Dmd*^Δ52^ rats, in which revertant fibres are rare. Long amplicon sequencing revealed that spontaneous exon skipping events lead to re-framing of the *Dmd* transcript and expression of a truncated dystrophin protein. The most prevalent transcript variant in *Dmd*^Δ45^ rats involved skipping of exon 44 (5.02% of total transcripts), followed by skipping of exons 46 and 47 (2.34%). In comparison, these events are nearly absent in WT rats (0.04% and 0.08%, respectively). These exon-skipping patterns mirror those observed in some DMD/BMD patients and may contribute to variability in clinical severity.

For instance, among 133 patients with an exon 45 deletion, 13 were reported to present with a milder, BMD-like phenotype [41]. In several of these cases, exon 44 skipping was identified as the mechanism allowing partial restoration of the reading frame and dystrophin expression [41]. Conversely, another patient with an exon 45 deletion exhibited minimal exon 44 skipping (∼1% of truncated *DMD* mRNA) and showed no signs of phenotype amelioration [43]. Such inter-individual variability—ranging from severe to mild phenotypes—is well documented in exon 45 deletions cases [15, 16, 20, 21] and is now recapitulated at the molecular level in this rat model.

Transcriptomic analysis of skeletal muscles in *Dmd^Δ45^*rats revealed both well-established and novel pathophysiological signatures. At 6 months, upregulation of genes related to regeneration and inflammation, such as S*ohlh2, Igfn1, Cdkn2a, Mymk*, and *Mymx*, suggests active myogenic repair and satellite cell engagement. However, elevated *Cdkn2a* expression also indicates potential cell cycle arrest, which may reflect satellite cell exhaustion or senescence. Genes involved in ECM remodelling and immune signalling—such as *Lrrc15*, *Tmem184a*, *Ptx4,* and *Clec2dl1—* were also upregulated, in line with ongoing fibrosis and chronic inflammation. These pathological responses were amplified at 12 months, with increased expression of these and other genes, indicating disease progression and intensification of compensatory pathways. Conversely, several genes were consistently downregulated at both 6 and 12 months, including *Barx2*, *Asb4*, *Ppp1r1a*, *RhoU*, and *Cacng7*, implicating disruption in myogenic signalling, ion transport and muscle structural maintenance. Additional downregulated genes at 12 months—such as *Unc5a*, *Slc6a2*, *Nppc*— point to worsening transcriptional repression as degeneration advances.

Overall, the transcriptional profile of skeletal muscle transitions from early stress response and regeneration at 6 months to chronic inflammation, fibrosis, and metabolic dysfunction by 12 months. Alongside established DMD markers (*Spp1*, *Thbs4*, *Postn*), novel gene candidates emerged, such as *Igfn1*, *Draxin, Cilp*, *Ctxn3* and *Fbln7* (upregulated), and *Barx2* (downregulated), offering new insights into disease mechanisms, biomarker discovery and therapeutic targets.

In the heart, transcriptomic profile revealed a temporal transition from early inflammatory and stress responses at 6 months to fibrotic remodelling and metabolic dysfunction at 12 months. At the earlier stage, Immediate Early Genes (IEGs) such as *Egr1*, *Fos*, *Nr4a1/2*, and *Arc* were upregulated, indicating rapid transcriptional responses to cellular and metabolic stress. Concurrently, inflammatory mediators (*Cxcl1*, *Socs3*, and *Nfkbiz)* and ECM remodelling genes (*Thbs1*, *Ccn1*, *Adamts1*) were also elevated, marking the onset of immune activation and tissue restructuring, which is consistent with a pre-fibrotic or early remodelling phase of cardiomyopathy in DMD. By 12 months, there was a widespread upregulation of fibrosis-related genes (e.g. *Comp*, *Fbln7*, *Fn1*, *Runx2, Abca1*), alongside downregulation of sarcomeric, ion channel, and metabolic genes (*Ryr1*, *Cacna1s*, *Pgam2)*, indicating contractile dysfunction, energy failure and advanced cardiac degeneration. Persistent up-regulation of metabolic regulators (*Fgf23*, *Abca1*, *Bcat1*) suggests ongoing metabolic stress in the degenerating cardiac tissue. This transcriptomic trajectory aligns with the natural course of DMD-associate cardiomyopathy, in which an early inflammatory phase evolves into irreversible fibrotic remodelling and functional defects.

In summary, this study successfully generated a *Dmd*^Δ45^ rat model that recapitulates the multisystemic features of DMD, including progressive skeletal muscle atrophy, cardiomyopathy, and neurobehavioral impairments. The presence of a significant proportion of revertant fibres likely contributes to the intermediate phenotype, differentiating it from more severe models like *Dmd^Δ52^*. The model offers a valuable platform for studying both the molecular basis of phenotypic variability and the mechanisms of spontaneous exon skipping. Our findings highlight exon 44 skipping as the predominant re-framing event in this model, suggesting that therapeutic strategies targeting exon 44 may be particularly effective in exon 45 deletion cases. Further investigation into the mechanisms regulating spontaneous exon skipping in *Dmd^Δ45^*rats could inform the development of personalized exon skipping therapies, with direct clinical relevance for a significant subset of DMD patients.

## MATERIAL AND METHODS

### *Dmd*^Δ45^ rat model generation

The *Dmd*Δ45 model was generated by Genoway (https://www.genoway.com/) by using one single guide RNA strategy (5’-TAGGAAGCTTGAGTCTGCGG-3’ with TGG PAM sequence) to target exon 45 in the rat Dystrophin gene (*Dmd*). The rats have a CD®(SD) (Crl:CD(SD)) background and were obtained from Charles River ((https://www.criver.com/fr). A 606bp deletion encompassing *Dmd* exon 45 **(Figure 1C)** and spanning from 207 bp into the 3’ region of intron 44 to 223 bp into the 5’ region of intron 45, including exon 45 was obtained. The *Dmd*^Δ45^ rat model line was back crossed at least 3 times with WT Sprague-Dawley rats received from Charles River. The rats were breed in a SPF barrier facility with 12h light/12 h dark cycles and were provided with food and water *ad libitum*. All rats were handled according to the European guidelines for the human care and use of experimental animals. All procedures on animals were approved by CERFE ethics committee C2EA-51 and the French Ministry of Research (MESRI) under the number APAFIS #25388.

### Genotyping

The genotyping of the rats was performed using the Phire Tissue Direct PCR Master Mix (Life technologies). Tail fragments extracted from rats were manipulated according to the supplier’s protocol. Samples were subjected to PCR with selected primers (**Table S7**).

### Histology

The muscle and heart tissues were frozen in isopentane and stored at -80°C. They were cut at the cryostat (Leica) and preserved at -80°C until staining. Transverse cryosections (8-10 µm) were prepared from frozen organs and processed for histological staining: hematoxylin-phloxine-saffron (HPS) and Sirius Red (SR) staining. Slides were scanned with the AxioScan (Zeiss) and stored at room temperature.

### Hematoxylin-phloxine-saffron (HPS) and Sirius Red (SR) staining

With the HPS stained muscle scans, QuPath (version 0.4.3) were used to train a small artificial neural network to classify positive and negative pixels for 3 categories: connective tissue; muscle tissue and inflammation. To assess tissue fibrosis, Sirius Red staining were used. For each muscle scan, QuPath (version 0.4.3) was used to classify positive and negative fibrotic pixels and subsequently used to quantify fibrotic areas.

### Tissue immunohistofluorescence and quantification of centronucleation and revertant fibres

Tissue sections were dried for 5 minutes and were rehydrated in PBS followed by blocking in either 10% Goat Serum or 10% FBS with 5% Goat Serum for 30 min. Primary and secondary antibodies **(Table S8)** were diluted in 1% Goat Serum (Agilent) or 1% FBS with 0.5% Goat The slides were scanned with AxiosScab (Zeiss) and stored at 4°C.

Fluorescence intensity, shape, and size were measured for each object (fibres, fibre membrane, and nuclei) using QuPath, an open-source software for bioimages analysis (version 0.3.2).. Centronucleation was estimated by measuring the distance between each nucleus and its closest membrane coordinate and normalized to the fibre’s minimal Feret diameter. Fibres with at least one internal nucleus located more than 1/5 of its minimal Feret diameter from the membrane were considered centronucleated. Positive fibres for any channel were detected based on the fluorescence distribution of negative control slices or slices from a known negative condition. DMD-positive fibres were identified by thresholding based on both absolute fluorescence intensity and relative intensity ratio between membrane and cytoplasm staining in a way to ensure 95% of positive fibres on WT rat samples.

### Western Blot

Harvested frozen muscle and heart tissues submerged in isopentane were lysed with RIPA buffer (ThermoFisher Scientific) containing EDTA-free protease inhibitor cocktail (Roche; 1 tablet per 10ml RIPA buffer) and 1/1000 diluted Benzonase (Sigma Aldrich). Primary and secondary antibodies were listed in **Table S8**. The membranes were revelated with the Odyssey 9120 (LI-COR) scanner. Signal was analysed with the program Image Studio Lite (LI-COR).

### RNA extraction and RT-PCR

Frozen muscle and heart tissues slices were lyzed in Nucleozol using the Fastprep. DNAse and RNAse free water was then added to the lyzed samples. RNA was extracted with Nucleomag extraction kit (Machery Nagel) according to the supplier’s protocol. Extracted RNA samples were treated for DNA removal with the TURBO DNA-free™ kit and Ribonuclease inhibitor RNasin (Promega) following the supplier’s protocol. Reverse transcription (RT) of mRNA was performed with the kit RevertAid H Minus First Strand cDNA synthesis (ThermoFisher Scientific) following the provider’s protocol. The obtained cDNA was then diluted to 1/4 using DNAse/RNAse-free water and amplified with a reverse transcription PCR reaction (RT-PCR) using Phusion® High-Fidelity DNA Polymerase (New England Biolabs) with different melting temperatures corresponding to the different primers used.

### RNA-seq analysis

Total RNA extraction was performed from muscle by the Trizol™ method (Thermo Fisher Scientific, Waltham, MA). Extracted RNA was dissolved in 20µl of RNase-free water and treated with Free DNA kit (Ambion) to remove residual DNA. Total RNA was quantified using a Nanodrop spectrophotometer (ND8000 Labtech, Wilmington Delaware).

RNA from 3 aliquots were sequenced with the sequencing depth between 18M and 50M reads per sample. RNA concentration was measured on a Nanodrop 2000 spectrophotometer (Thermo Fisher Scientific). RNA quality (RIN≥7) was controlled using an Agilent RNA 6000 Pico Kit on a 2100 Bioanalyzer instrument (Agilent Technologies). The sequencing libraries were prepared using the TruSeq Stranded Total RNA Library Prep Kit (Illumina) and sequenced according to the Illumina protocol. Paired-end reads (2x150bp) were aligned on the reference transcriptome (Rattus_norvegicus. mRatBN7.2) using STAR version 2.7.11b [45] with an average alignment of 95.0%, after excluding samples showing at the same time a low number of reads and a low percentage of alignment. Gene expression was measured by featureCounts version 2.0.6. Quantification files were processed using R (4.4.1) to perform differential expression analysis. Pairwise group differential expression were performed using DESeq2 (1.44.0) considering genes with more than 50 reads per samples (lfcThreshold = 0, pAdjustMethod = “fdr”, independentFiltering = T). Genes were considered dysregulated if the absolute Log2 fold change was above 1 and the FDR adjusted p.value below 0.05.

### Seric biomarker quantification

CK activity was measured by colorimetry using the Dri-chem (Fujifilm). Serum was diluted in MilliQ water and deposited onto FUJI DRI-CHEM Slides for CPK-PIIIS/Creatine Phospho Kinase (DMV imaging).

Enzyme-Linked Immunosorbent Assay (ELISA) was performed on serum to quantify MYOM3 using a Meso Scale Discovery assay (MSD) according to a previously described protocol [46]. Briefly, the multi-array plate containing electrodes (MSD) was first coated overnight at 4°C with the capturing antibody for MYOM3 (17692-1-AP, Proteintech). After washes with PBS + 0.05% Tween 20, a saturation of the plate was performed by adding 3% bovine serum albumin (BSA) 1h at RT. New washes were performed followed by the incubation of the diluted sera (in 1% BSA solution, 2h at RT with agitation). To calculate the concentration of the protein, a standard range of a MYOM3 peptide corresponding to the antibody epitope (synthesized by Proteogenix) was used. The antibody for detection was an in-house anti MYOM3 (Clone 51-H1-B4, subcontracted to Proteogenix©) coupled with sulfo/TAG (Mesoscale).

### Isolated muscle force measurement

Animals were sacrificed by intraperitoneal injection of a lethal dose of Dolethal. The muscles (EDL and TA) were then dissected and soaked in an oxygenated Tyrode solution (95% O2 and 5% CO2) maintained at a temperature of 20°C. Muscles were connected at one end to an electromagnetic puller and at the other end to a force transducer. Stimulation was delivered through electrodes running parallel to the muscle.

Twitch and tetanic (rat EDL: 125Hz, 500ms; Sol: 2400 ms, 90Hz) isometric contractions were studied at Lo (the length at which maximal tetanic isometric force is observed). Each stimulation was performed at 600 mA. For comparative purposes, normalized isometric force (or tension) instead of force was assessed. Isometric tension was calculated by dividing the force by the estimated cross-sectional area (CSA) of the muscle. Assuming muscles have a cylindrical shape and a density of 1.06 mg.mm-3, CSA corresponds to the wet weight of the muscle divided by its fibre length.

### Electromyography

Electromyographic recording allows neurophysiological measurement of motor and sensory function. The rats were anaesthetized intraperitoneally (injection volume = 2 ml/kg; ketamine concentration =100 mg/kg; xylazine concentration = 10 mg/kg). This test was performed with an EMG device (NATUS). Needle electrodes (30G) were used to stimulate the nerve and receive the electrical response at the muscle. The needles were inserted through the animal’s skin to a depth of 1-2 mm. Four trains of 200 stimuli (0.1 ms, 8 mA) at a frequency of 10 Hz were applied to the sciatic nerve with an interval of 1 second between two trains. The amplitude of the CMAP at the level of the gastrocnemius muscle was measured 30 sec before and 30 sec after the four stimulus trains with a single stimulus (8 mA, 0.1 ms). The percentage decrease in the amplitude of the muscle response between the response before and after the four stimulus trains is used as a measure of the level of muscle fatigue.

### Echocardiography

Conventional echocardiography was performed on anesthetized mice using a Vevo 3100LT (Visual sonics) with a 40 MHz mice cardiac probe (MX550D) or 15 MHz rats cardiac probe (MX201). During the procedure, heart rate (HR) and temperature were monitored. For echocardiography recording, sweep speed, depth, focus and gain settings were optimized to obtain the most defined acquisitions. Two-D and M-mode echocardiography were performed manually from the long parasternal axis view at the level of the largest LV diameter (at the level of the papillary level). LV dimensions [LV end-diastolic diameter (LVEDD), LV end-systolic diameter (LVESD), posterior wall (LVPW) and anterior wall (LVAW) wall thickness] were measured using the leading-edge convention of the American Society of Echocardiography. The shortening fraction (SF) and EF of the left ventricle and LV mass were calculated from the above dimensions. Pulmonary and aortic artery velocity and pressures to detect intra-cardiac pressures changes (Aov and RV function) was evaluated using color Doppler imaging in pulmonary artery and aortic arch.

### Behaviour study on open field

Rats were tested in automated 90*90*40 cm open fields (Panlab, Barcelona, Spain), each virtually divided into central and peripheral regions. The open field was placed in a room homogeneously illuminated at 15 Lux. Each rat was placed in the periphery of the open field and allowed to explore freely the apparatus for 30 min, with the experimenter out of the animal’s sight. The distance travelled, the number of rearings, and time spent in the central and peripheral regions were recorded over the test session (peripheral width = 20 cm). The number of entries and the percentage of time spent in centre area were used as index of emotionality/anxiety.

### Y maze spontaneous alternation

Testing occurs in a Y-shaped maze with three white, opaque plastic arms (90*25*40 cm) at a 120° angle from each other. Each arm has specific patterns on the walls. After introduction to the center of the maze, the animal was allowed to freely explore the three arms during 8 min. Over the course of multiple arm entries, the subject should show a tendency to enter a less recently visited arm. The number of arm entries and the number of triads were recorded in order to calculate the percentage of alternation. An entry occurs when all four limbs are within the arm.

### Statistics

The statistical analyses were performed after verification of the normal distribution of the data with the Shapiro-Wilk test. If the data followed a normal distribution, grouped data were analysed by two-way ANOVA, and non-grouped data were analysed by t test. Data that did not follow a normal distribution were analysed by the non-parametric test Kruskal-Wallis. The results of statistical analysis were represented with ns: non-significant; *: p<0.05; **: p<0.01; ***: p<0.001; ****: p<0.0001.

## ACKNOWLEDGMENTS

This work was supported by the “Association Française contre les Myopathies” (AFM) (Funding n° 23853), and “Institut National de la Santé Et de la Recherche Médicale”. The authors are Genopole’s members, first French biocluster dedicated to genetic, biotechnologies and biotherapies. We are grateful to the “Imaging and Cytometry Core Facility” and to the *in vivo* evaluation services of Genethon for technical support, to Ile-de-France Region, to Conseil Départemental de l’Essonne (ASTRE), INSERM and GIP Genopole, Evry for the purchase of the equipment. We thank Célia Thevenard for her help in figure design and Lydie Debaize for comments on the manuscript.

## COMPETING INTERESTS

The author(s) declared no potential conflicts of interest with respect to the research, authorship, and/or publication of this article.

## AUTHOR CONTRIBUTIONS

I.R. was responsible for experimental design and project management. A.J., M.B. and G.W. helped supervise the project. C.D., J.B., A.Do., A.Du, and L.P. performed experiments. T.W. managed the RNA-seq experiment and analysis. G.C assisted with RNA-seq data processing and analysis. S.A managed the functional analysis of the DMD rat. A.J., T.W. and I.R. wrote the manuscript with input of all authors.

---

**Figure S1.**
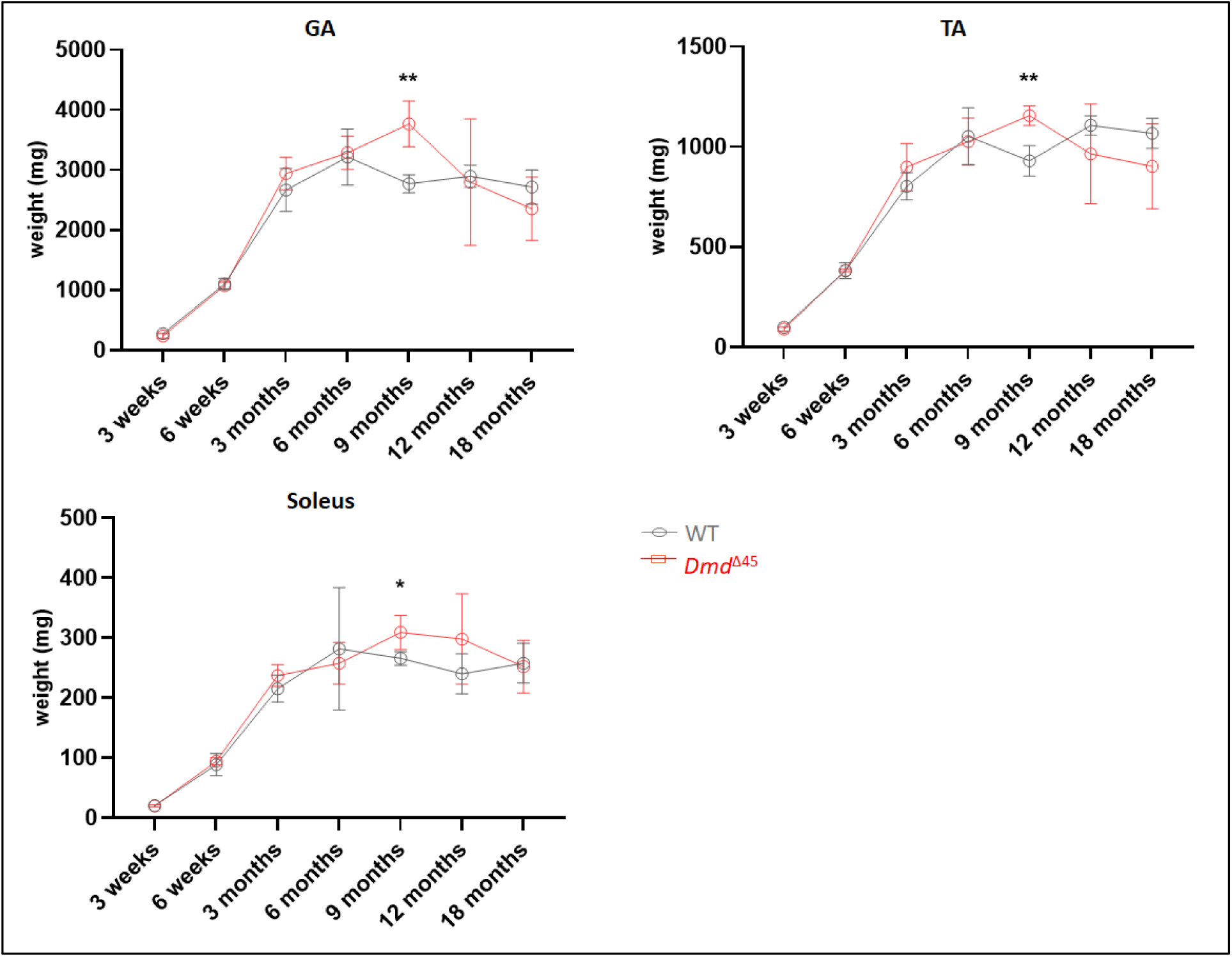
Muscle weight measurement of WT and *Dmd^Δ45^* rats from 3 weeks and 18 months (n=3-10).

**Figure S2:**
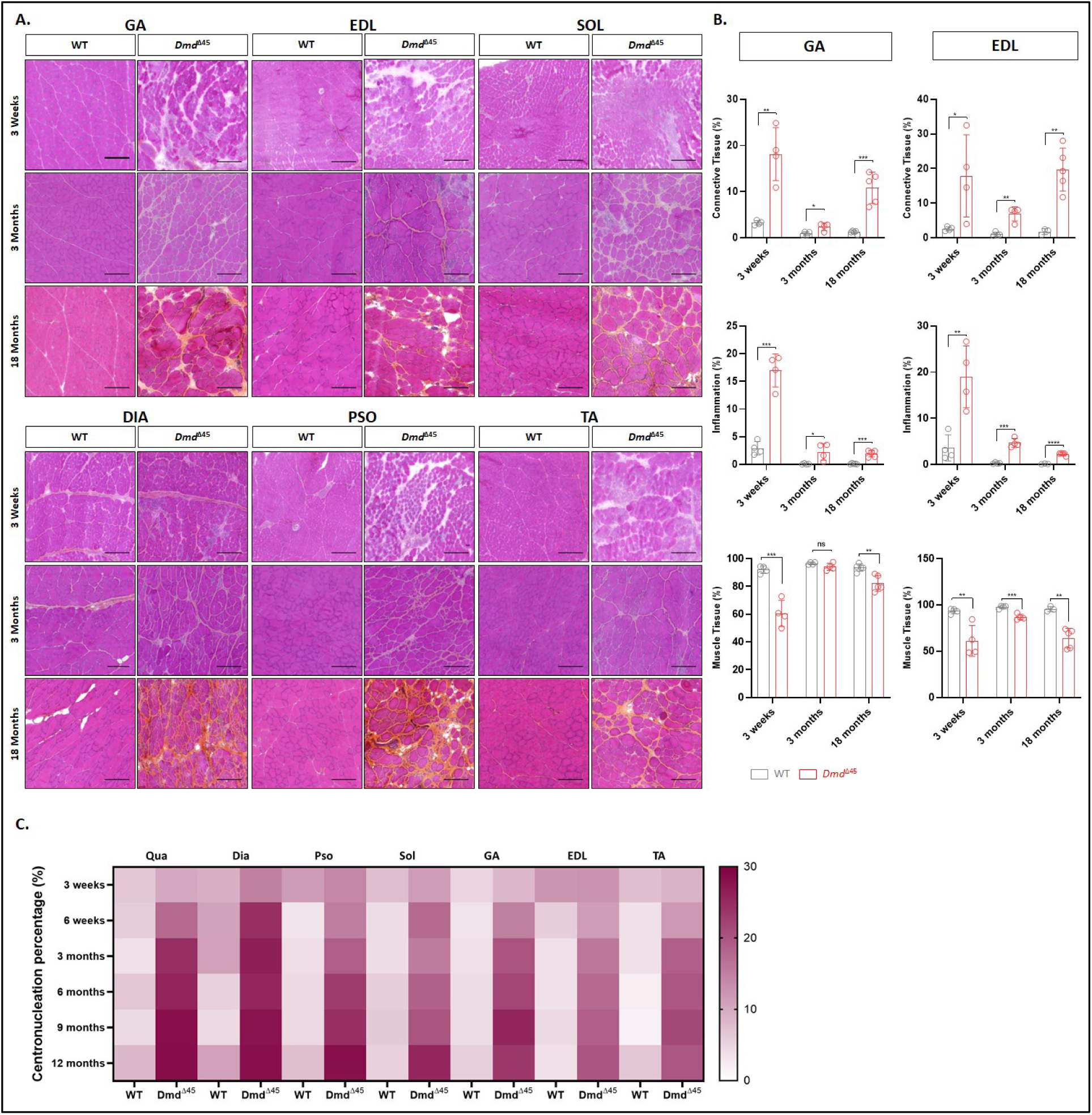
Histological evaluation of skeletal muscles. **A)** Representative images of skeletal muscles, Gastrocnemius (GA), EDL, Soleus (SOL), Diaphragm (DIA), Psoas (PSO) and tibialis anterior (TA) cross-sections stained with Hematoxylin Phloxine Saffron (HPS) from WT and *Dmd^Δ45^*rats at 3 weeks, 3 months and 18 months. Size bar = 200 µm. **B)** Quantification of connective tissue, inflammation and muscle tissue percentages from HPS-stained cross-sections in GA and EDL (n = 3–5). **C)** Heatmap showing centronucleation percentage in different skeletal muscles of WT and *Dmd^Δ45^* across ages.

**Figure S3:**
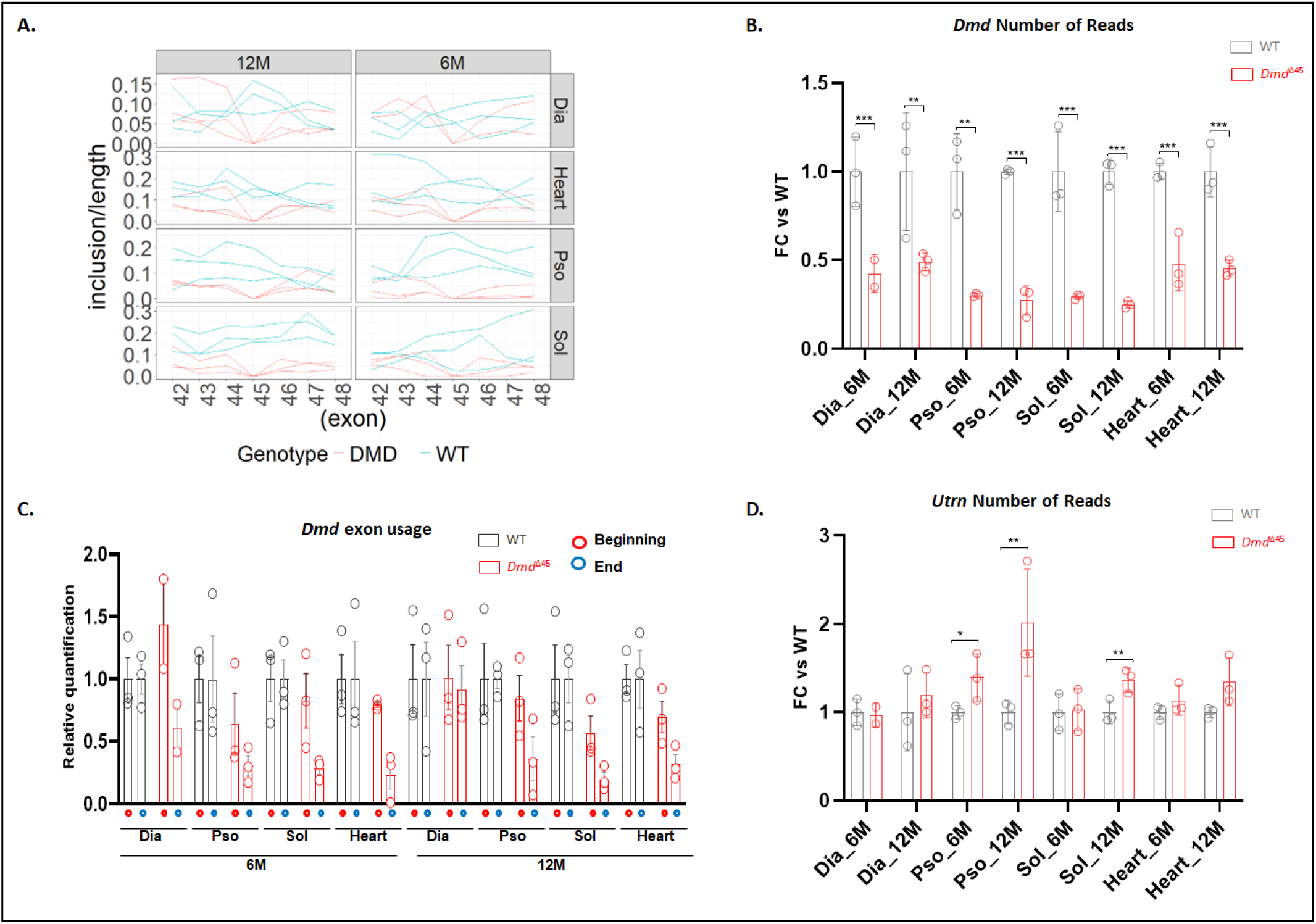
RNA-seq analysis revealed dysregulated pathways associated to DMD. **(A)** Relative read counts for exons 42 to 48 of the *Dmd* gene in Dia, Pso, Sol, and heart tissues at 6 and 12M. **(B)** Relative read counts for *Dmd* in Dia, Pso, Sol and Heart samples at 6M or 12M. **(C)** Relative total exon read counts for *Dmd* in Dia, Pso, Sol and Heart samples at 6M or 12M. The Exons were grouped into two regions: Beginning (exons upstream of exon 45) and End (exon downstream of exon 45). **(D)** Relative read counts of Utrophin (*Utrn*) of the Dia, Pso, Sol and Heart samples at 6M or 12M.

**Figure S4:**
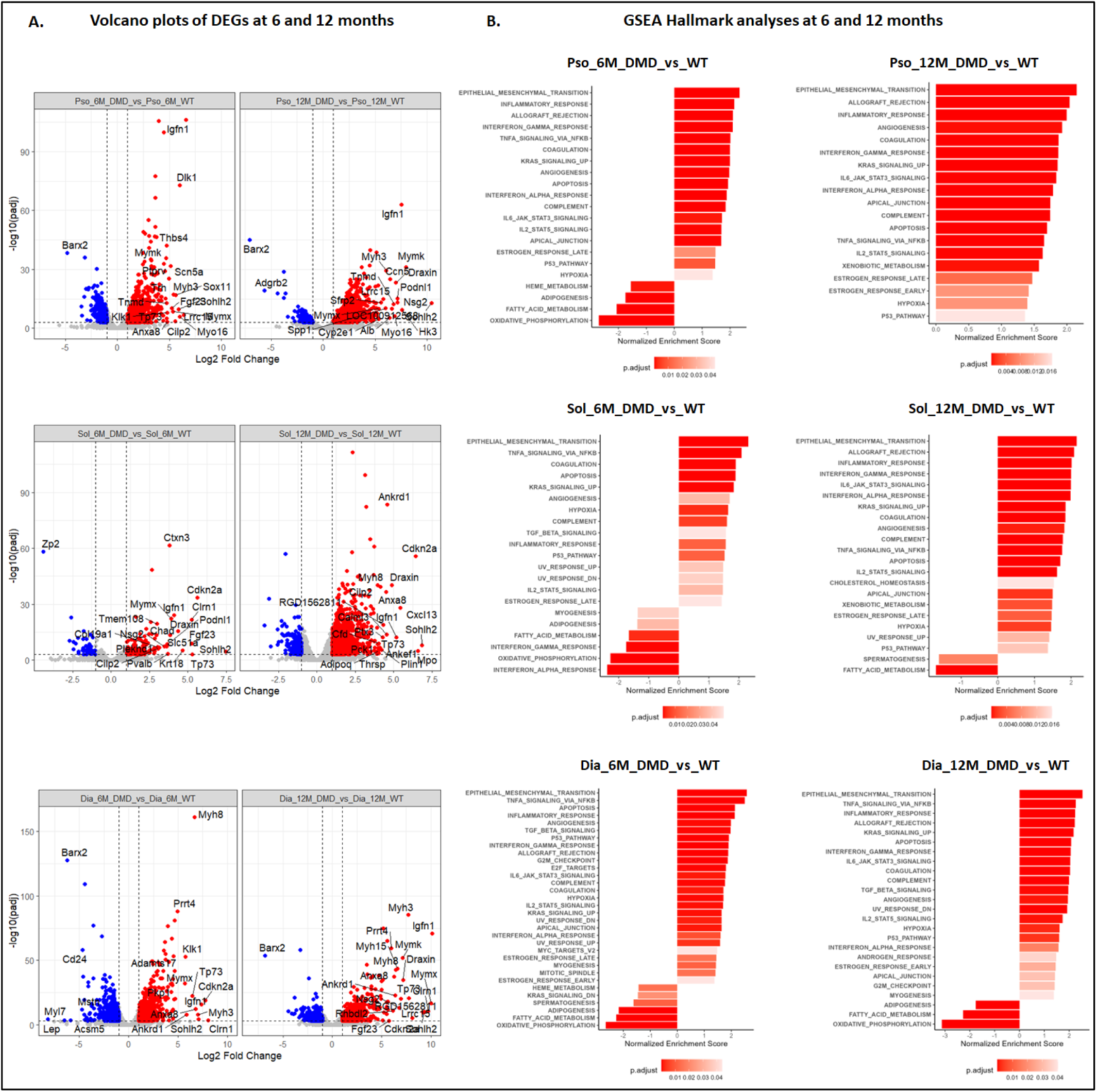
**(A**) Volcano plots showing differential gene expression in Sol and Dia muscle samples, comparing *Dmd^Δ45^* and WT rats at 6M and 12M. **(B)** Bar plot showing the GSEA Hallmark pathway analysis in skeletal muscles.

**Figure S5:**
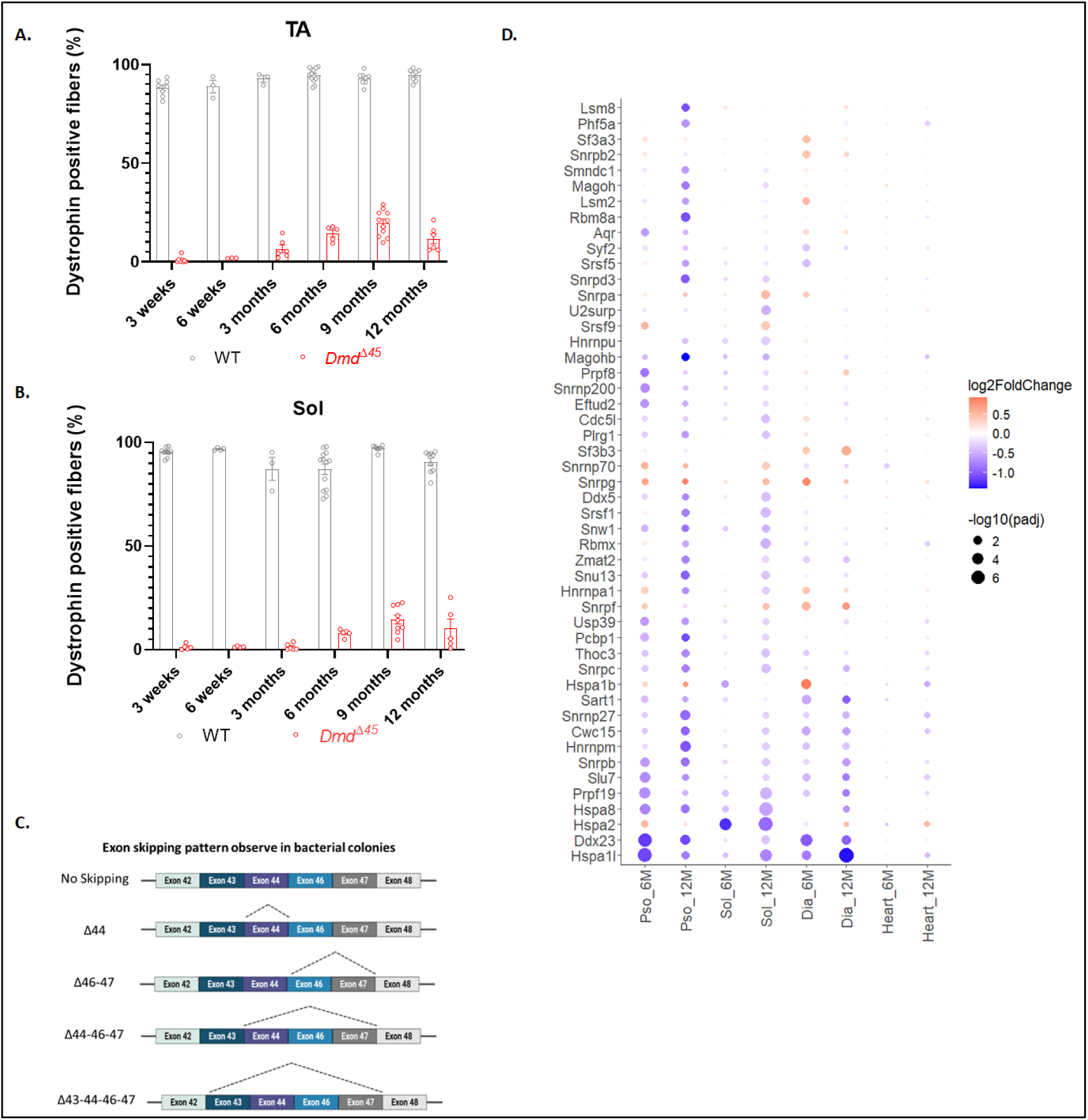
Partial restoration of dystrophin expression and dysregulation of spliceosome related genes in Dmd^Δ45^ rat model. A-B) Quantification of the number of dystrophin positive fibres in different muscles from 3 weeks to 12 months of age. **C)** Analysis of exon skipping event showing the exon skipping pattern. **D)** Dot graph of the dysregulated spliceosome related genes. Each dot represents a gene in a specific DMD sample. Colour indicates the log₂ fold change, with blue for downregulation and red for upregulation. Dot size reflects the -log10(padj) for that gene in the corresponding DMD sample. Genes are ordered based on the geometric mean of their adjusted p-values across all conditions.

**Table S1:**
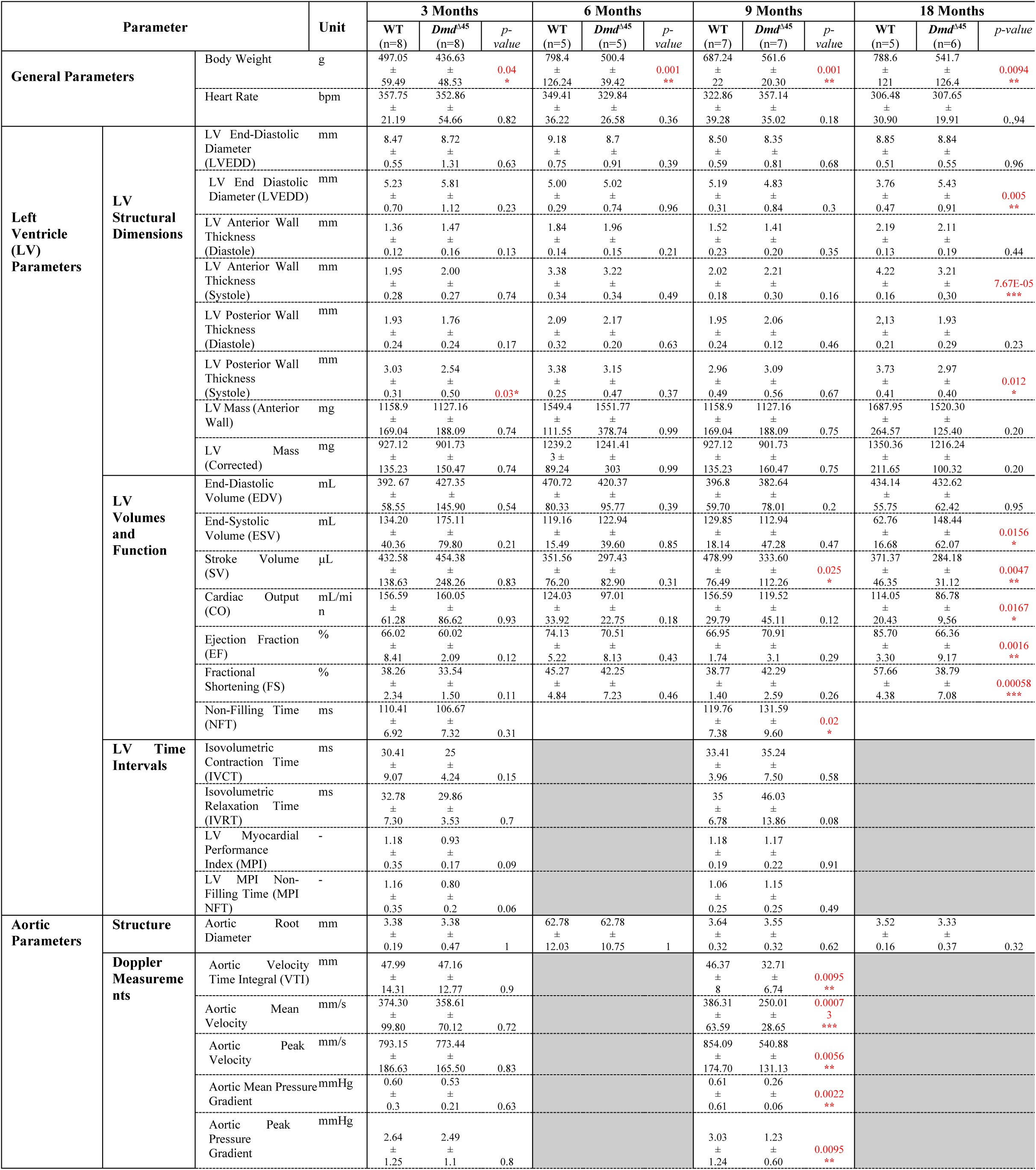
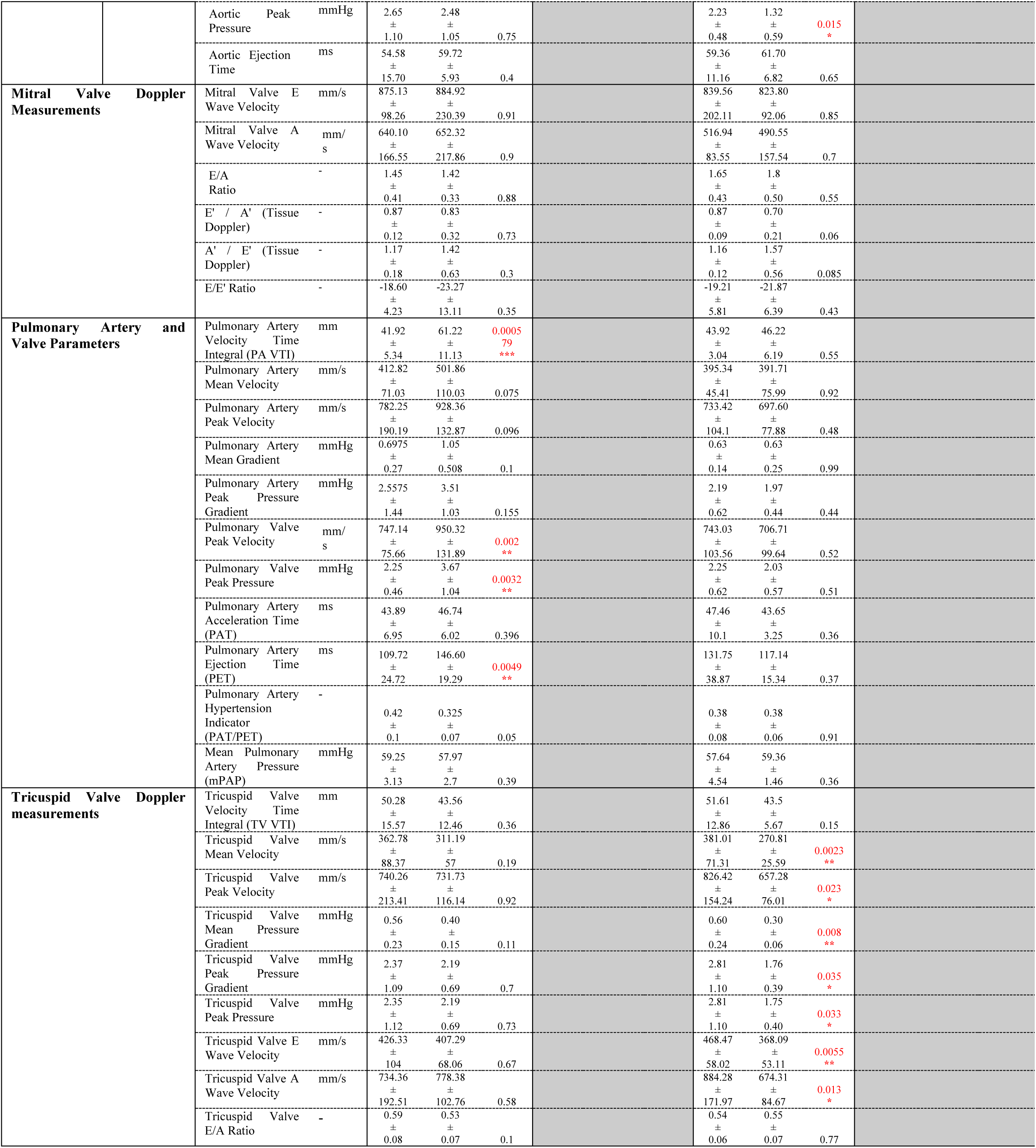
Transthoracic echocardiography parameters. Doppler image analyses were performed only in 3- and 18 months old rat. Significant p-values are indicated in red.

**Table S2:** Top contributing genes to PC1. Genes with an average contribution score greater than 0.03 from PCA were selected and ranked by absolute contribution score. References where the dysregulation of the corresponding genes was reported are indicated.

**Upregulated genes**
| gene | Full name | Ref in DMD |
| --- | --- | --- |
| Ankrd1 | Ankyrin Repeat Domain 1 | [48] |
| Igfn1 | Ig Like And Fibronectin Type III Domain Containing 1 | - |
| Myh3 | Myosin Heavy Chain 3 | [49] |
| Myh8 | Myosin Heavy Chain 8 | [50] |
| Thbs4 | Thrombospondin 4 | [51] |
| Cdkn1a | Cyclin Dependent Kinase Inhibitor 1A | [52] |
| Comp | Cartilage Oligomeric Matrix Protein | [27] |
| S100a4 | S100 Calcium Binding Protein A4 | - |
| Cilp | Cartilage Intermediate Layer Protein | - |
| Anxa8 | Annexin A8 | - |
| Postn | Periostin | [53] |
| Dclk1 | Doublecortin Like Kinase 1 | - |
| Cdkn2a | Cyclin Dependent Kinase Inhibitor 2A | [54] |
| Mt2A | Metallothionein 2A | - |
| Draxin | Dorsal Inhibitory Axon Guidance Protein | - |
| Mymk | Myomaker, Myoblast Fusion Factor | [55] |
| Prrt4 | Proline Rich Transmembrane Protein 4 | - |
| Ctxn3 | Cortexin 3 | - |
| Timp1 | TIMP Metalloproteinase Inhibitor 1 | [56] |
| Mymx | Myomixer, Myoblast Fusion Factor | [55] |
| Gpnmb | Glycoprotein Nmb | [39] |
| Prrg4 | Proline Rich And Gla Domain 4 | - |
| Tubb6 | Tubulin Beta 6 Class V | [57] |
| Spp1 | Secreted Phosphoprotein 1/ Osteopontin | [58] |
| Myl4 | Myosin Light Chain 4 | [51] |
| Klk1 | Kallikrein 1 | - |
| Mt1a | Metallothionein 1A | - |
| Tmem158 | Transmembrane Protein 158 | - |
| Tnnt2 | Troponin T2, Cardiac Type | [59, 60] |
| Fbln7 | Fibulin 7 | - |
| Cd24 | CD24 Molecule | - |
| Col11a1 | Collagen Type XI Alpha 1 Chain | - |
| Socs3 | Suppressor of cytokine signalling 3 | - |

**Downregulated genes**
| gene | Full name | Ref in DMD |
| --- | --- | --- |
| Barx2 | BARX Homeobox 2 | - |
| Mylk4 | Myosin Light Chain Kinase Family Member 4 | - |
| Aqp4 | Aquaporin 4 | [61] |

**Table S3:** Commonly dysregulated genes among skeletal muscles. The genes are ranked by the average log₂ fold change (FC) across samples. The 20 genes with the highest average fold change were selected out of 26 genes for down-regulated genes at 6 months, 179 genes for up-regulated genes at 6 months, 58 genes for down-regulated genes at 6 months and 625 genes for up-regulated genes at 12 months. Genes that are found at the 2 time points are in red. Genes that are found in the top contribution PC1 list are highlighted in yellow. The DMD gene is in bold.

| Up_6M | FC | Up_12M | FC | Down_6M | FC | Down_12M | FC |
| --- | --- | --- | --- | --- | --- | --- | --- |
| **Sohlh2** | 6.80 | **Sohlh2** | 8.84 | **Barx2** | -4.47 | **Barx2** | -5.41 |
| **Igfn1** | 6.03 | **Igfn1** | 7.57 | Zp2 | -3.23 | Unc5a | -3.53 |
| Clrn1 | 5.76 | **Draxin** | 6.33 | Oxtr | -3.16 | **AABR07021734.1** | -3.51 |
| **Cdkn2a** | 5.69 | **Lrrc15** | 6.20 | **AABR07021734.1** | -2.96 | Slc6a2 | -3.07 |
| Pilrb-ps6 | 5.63 | Th | 6.20 | **Asb4** | -2.69 | **Asb4** | -3.05 |
| Serpina3c | 5.48 | **Mymk** | 6.17 | Adra2c | -2.48 | Nppc | -2.88 |
| **Tp73** | 5.48 | **Cdkn2a** | 6.12 | Adig | -2.43 | Gm37419 | -2.41 |
| **Mymx** | 4.98 | **Clec2dl1** | 6.02 | Dpep1 | -2.18 | Mogat2 | -2.33 |
| **Fgf23** | 4.94 | **Nsg2** | 5.84 | **Ppp1r1a** | -2.13 | **Cacng7** | -2.32 |
| **Anxa8** | 4.78 | Bin2a | 5.59 | **Rhou** | -2.08 | LOC102551557 | -2.26 |
| **Klk1** | 4.60 | LOC120103148 | 5.57 | **Cacng7** | -2.03 | Gucy2g | -1.98 |
| Gbx2 | 4.58 | Tnn | 5.57 | Inpp5j | -1.84 | Rnf43 | -1.97 |
| Serpinb2 | 4.49 | Otog | 5.51 | Fbxo44 | -1.82 | **Ppp1r1a** | -1.94 |
| **Ankrd1** | 4.46 | Klra5 | 5.38 | Smtnl2 | -1.69 | Spdya | -1.94 |
| **Lrrc15** | 4.37 | **Ptx4** | 5.21 | PCOLCE2 | -1.62 | **Rhou** | -1.92 |
| **Tmem184a** | 4.23 | Bicdl2 | 5.18 | LOC120098897 | -1.61 | P3r3urf | -1.86 |
| **Draxin** | 4.22 | **Podnl1** | 5.13 | **Dmd** | -1.59 | Nrep | -1.84 |
| **Ptx4** | 4.22 | **RGD1562811** | 5.09 | Trim45 | -1.58 | Pcdhga1 | -1.73 |
| **Rassf7** | 4.21 | Tnmd | 5.07 | Ppm1j | -1.56 | AABR07052523.2 | -1.71 |
| **Clec2dl1** | 4.16 | **Tmem184a** | 5.04 | Rab3b | -1.47 | Tuba8 | -1.69 |

**Table S4:**
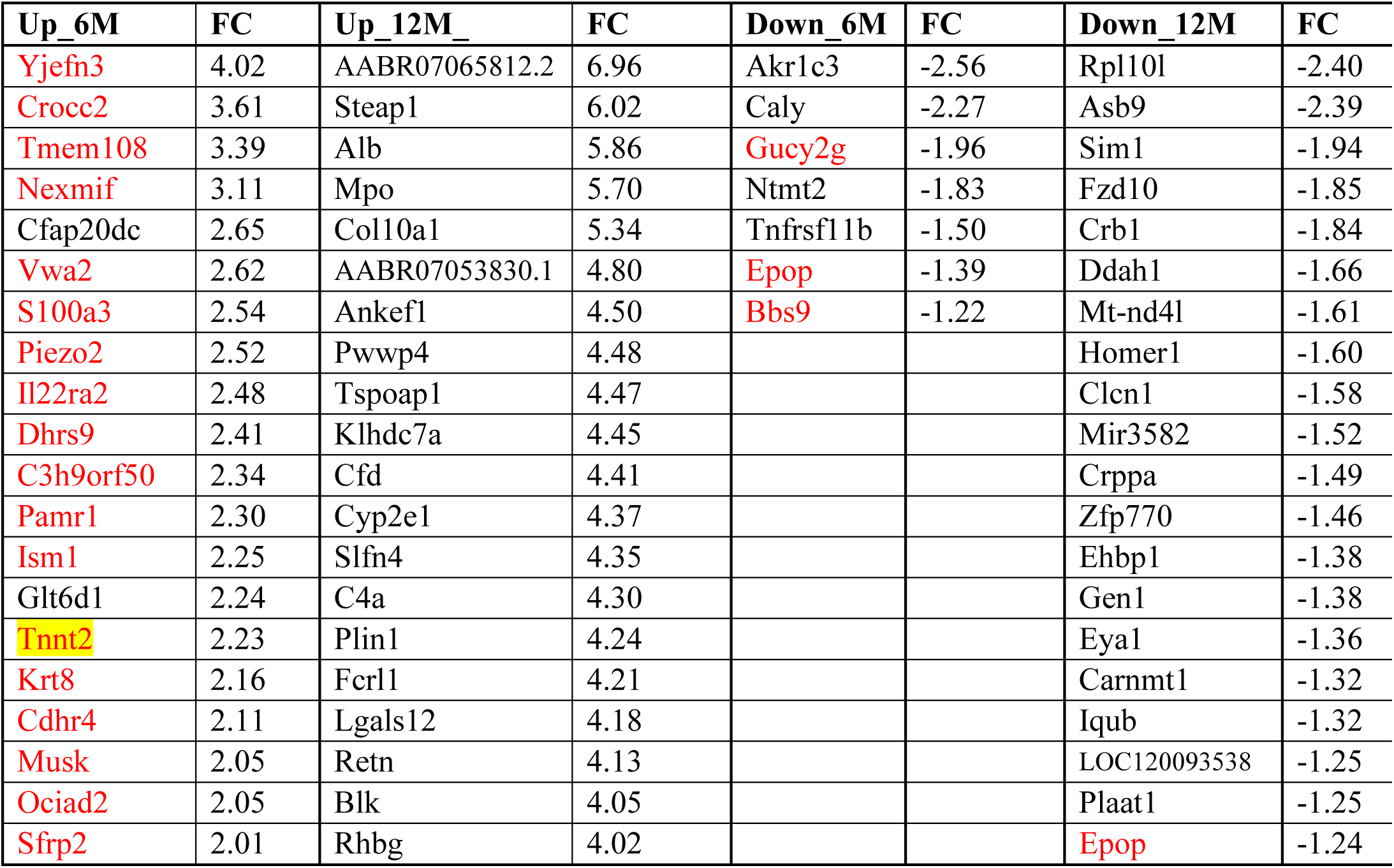
Additional dysregulated genes in both Pso and Sol. The genes are ranked by the average log₂ fold change across samples. When more than 20 genes were identified, those with the highest average fold change were selected with 20 out of 34 for the down regulated genes at 6M, 20 out of 56 for the upregulated genes at 6 months and 20 out of 797 at 12 months. Genes that are found at the 2 time points are in red. Genes that are found in the top contribution PC1 list are highlighted in yellow.

**Table S5:** Commonly dysregulated genes in heart. The genes are ranked by the log₂ fold change (FC) across samples. Only the top 20 first genes are shown. Genes that are found at the 2 time points are in red. The *dmd* gene is in bold.

| 6M UP |  | 12M UP |  | 6M DN |  | 12M DN |  |
| --- | --- | --- | --- | --- | --- | --- | --- |
| Gene | FC | Gene | FC | Gene | FC | Gene | FC |
| Pilrb-ps6 | 4.41 | Spp1 | 3.81 | LOC100912568 | -7.47 | Tnnc2 | -4.60 |
| Fgf23 | 4.31 | Kcne1 | 3.60 | **Cacna1s** | -3.67 | **Cacna1s** | -3.97 |
| Cxcl2 | 3.28 | Clec2dl1 | 3.36 | AABR07021734.1 | -3.63 | Actn3 | -3.86 |
| Egr1 | 3.25 | Mmp12 | 3.15 | Adh6 | -3.35 | Atp2a1 | -3.78 |
| Hist1h4m | 3.09 | Thbs4 | 3.12 | Zp2 | -3.19 | Pvalb | -3.41 |
| Fos | 3.00 | Ltbp2 | 2.84 | **Slc17a7** | -2.65 | Tnni2 | -3.08 |
| Nr4a1 | 2.44 | Olr1 | 2.75 | RT1-CE16 | -2.47 | Slc4a1 | -3.03 |
| Cxcl1 | 2.37 | Trh | 2.65 | Olr35 | -2.39 | Myh2 | -2.90 |
| Egr3 | 2.35 | **Thbs1** | 2.61 | Rpl39 | -2.10 | Smtnl1 | -2.84 |
| Btg2 | 2.10 | Nefm | 2.58 | Lrrn2 | -1.82 | Myl11 | -2.81 |
| Arc | 2.08 | **Rgs1** | 2.43 | Fam222a | -1.72 | Mylk2 | -2.73 |
| **Egr2** | 2.06 | Serpina3a | 2.36 | Neb | -1.67 | Ryr1 | -2.58 |
| Ier2 | 2.03 | Nrg1 | 2.28 | Alpg | -1.55 | Slc9a2 | -2.32 |
| Zfp36 | 1.91 | Ptgs2 | 2.28 | **Ano5** | -1.41 | **Fam131c** | -2.09 |
| Rgs1 | 1.84 | Comp | 2.27 | Pxmp4 | -1.41 | **Slc17a7** | -2.04 |
| Ccn1 | 1.80 | Fhad1 | 2.26 | **Dmd** | -1.35 | Myom3 | -1.90 |
| Pygl | 1.74 | Tnfrsf11b | 2.18 | **Csdc2** | -1.34 | Atp6ap11 | -1.79 |
| Nr4a2 | 1.67 | Piezo2 | 2.16 | AY172581.19 | -1.33 | Pnpla3 | -1.75 |
| **Thbs1** | 1.62 | Crtac1 | 2.15 | Atp1a3 | -1.28 | H2ac1 | -1.73 |
| Nfkbiz | 1.51 | Olfm2 | 2.14 | Susd5 | -1.25 | Crybb1 | -1.69 |

**Table S6:**
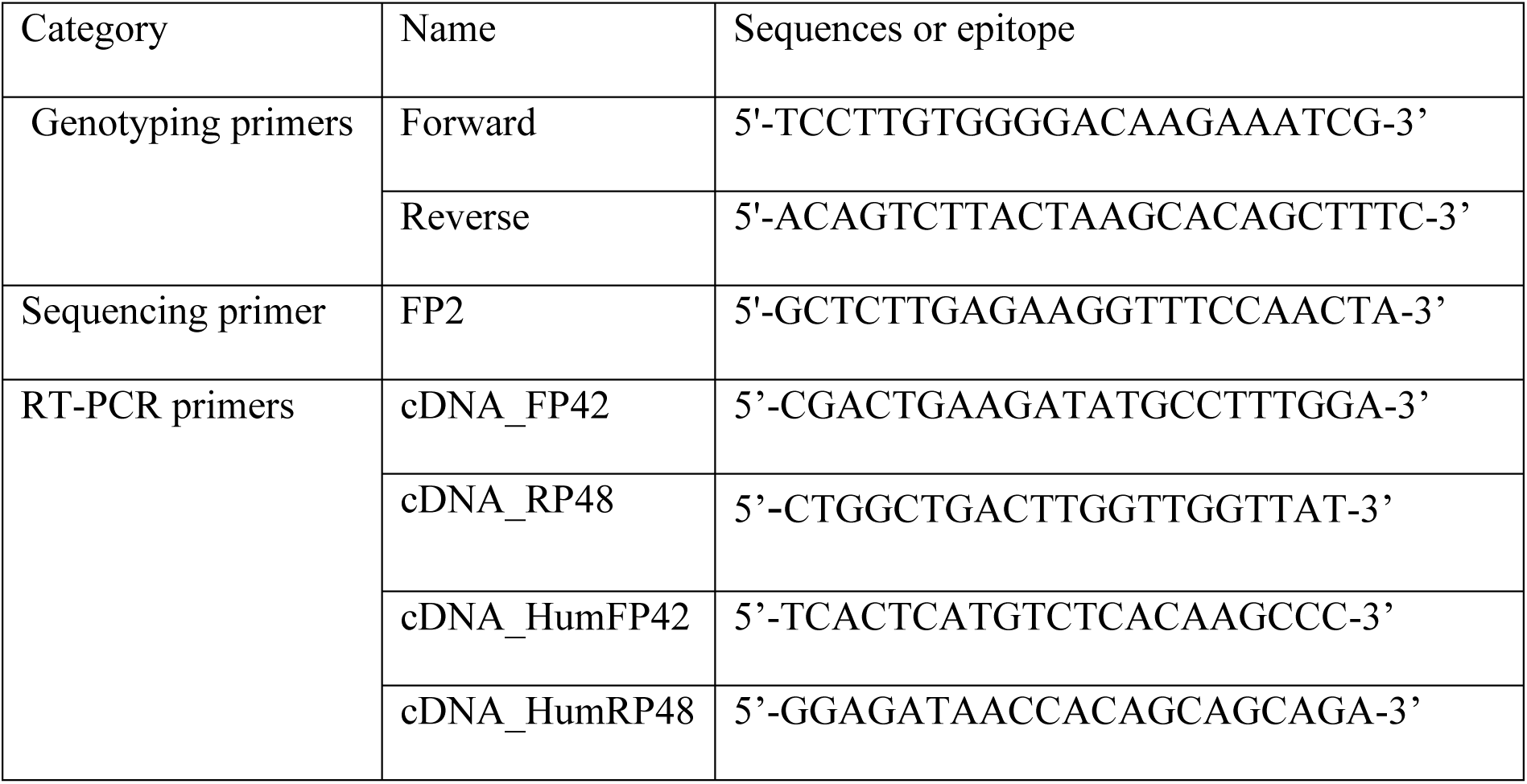
Primer sequences.

**Table S7:** Antibodies.

**A) Immunohistofluorescence antibodies**
| **Name** | **Recognised epitope** |
| --- | --- |
| Dys-b (Leica *NCL-DYSB*) | Dystrophin N-terminal |
| Dys-2 (Leica *NCL-DYS2*) | Dystrophin C-terminal |
| Anti-dystrophin (Abcam *ab15277*) | Dystrophin C-terminal |
| Anti-laminin (Abcam *ab11575*) | Laminin |
| Anti-utrophin (Leica *DRP2-CE*) | Utrophin N-terminal |
| Anti-MHCd (Leica *NCL-MHCd*) | MHCd |
| Alexa fluor 594 Donkey anti-rat IgG Highly Cross-Adsorbed Secondary (Life technologies *A-21209*) | Rat IgG (Heavy and light chains) |
| Alexa fluor 594 Donkey anti-mouse (Life technologies *A21203*) | Mouse IgG (heavy and light chains) |
| Alexa fluor 488 Goat anti-rabbit (Life technologies *A11008*) | Rabbit IgG (heavy and light chains) |

**B) Western Blot antibodies**
| **Name** | **Recognized epitope** |
| --- | --- |
| Dys-b (Leica *NCL-DYSB*) | Dystrophin N-terminal |
| Dys-2 (Leica *NCL-DYS2*) | Dystrophin C-terminal |
| Anti-alpha actinin 4 (Life technologies *PA5-22259*) | Alpha actinin 4 (within amino acids 206 and 542) |
| IRDye® 800CW Donkey anti-Mouse IgG Secondary Antibody (Li-cor *926-32212*) | Mouse IgG (Heavy and light chains) |
| IRDye® 680CW Donkey anti-Rabbit IgG Secondary Antibody (Li-cor *926-68073*) | Rabbit IgG (Heavy and light chains) |

## Notes

### Competing Interest Statement

The authors have declared no competing interest.

